# Uncertainty-aware breeding decisions: MCMC-based optimum contribution selection increases breeding decision robustness

**DOI:** 10.64898/2026.03.15.711440

**Authors:** Jon Ahlinder, Patrik Waldmann

## Abstract

Optimum contribution selection (OCS) balances genetic gain and inbreeding by optimizing parental contributions to the next generation, but current implementations rely on point estimates of breeding values that discard the uncertainty inherent in genetic evaluations. We introduce CVaR-OCS, a novel formulation that incorporates the full posterior distribution of estimated breeding values (EBVs) directly into the OCS objective via Conditional Value at Risk (CVaR) (a coherent risk measure from financial portfolio theory) allowing a single optimization to simultaneously maximize expected genetic gain and protect against worst-case outcomes driven by EBV uncertainty. We evaluate CVaR-OCS on a simulated multi-generation genomic selection dataset with known true breeding values (QTL-MAS 2010; *n* = 900 candidates), enabling direct comparison to an oracle solution, and on Norway spruce (*Picea abies n* = 5,525) forest tree breeding progeny trials. On the simulated dataset, MAP-OCS overestimated its own expected genetic gain by 16.7% due to EBV uncertainty, a bias eliminated by CVaR-OCS by construction, while CVaR-OCS also recovered one additional oracle-optimal individual and reduced gain distribution variance by 4.8%. In Norway spruce, the recommended CVaR-OCS operating point improved tail-gain security by 6.60% and broadened the selection base from 145 to 159 individuals at a genetic gain cost of only 0.70%. Complementary MCMC-based robustness scores revealed that 25 MAP-OCS selections in Norway spruce were unstable across the posterior distribution; post-hoc exclusion of these individuals failed to improve tail-gain security, motivating the principled CVaR-OCS approach. CVaR-OCS provides breeders with a principled, computationally efficient tool for uncertainty-aware selection decisions, and multi-generation simulation studies are needed to fully characterize its long-term effects on genetic gain trajectories and inbreeding accumulation.

## Introduction

Maximizing genetic merit while restricting relatedness in a breeding population is a major challenge in many breeding programs (Lynch and Walsh 1998; Goddard 1994). If relatedness is not controlled, elevated levels of inbreeding will cause fitness depression and reduce the performance of the offspring in the population (Lynch and Walsh 1998; Charlesworth and Willis 2009). Typically, the genetic load will increase, that is, the presence of deleterious alleles in the population, leading to a higher incidence of genetic disorders (Derks and Steensma 2021; Tao *et al*. 2025). Therefore, much effort has been devoted to developing methods that optimize the balance between gain and relatedness of the selected set of individuals and their contribution to the next generation of breeding or deployment population (Goddard 1994; Mullin and Belotti 2015; Hamilton 2020; Cowling *et al*. 2023; Tchounke *et al*. 2023).

One of the most popular approaches, the optimum contribution selection (OCS), optimizes a linear objective function with quadratic constraints on relatedness (inbreeding) (Meuwissen 1997). To solve the optimization problem, the original approach uses Lagrangian multipliers in an iterative fashion where non-selected individuals are excluded one by one. Since then, many efforts have been made to further improve the original approach by introducing enhanced optimization techniques such as semidefinite programming (SDP) (Pong-Wong and Woolliams 2007), the branch-and-bound algorithm (Mullin and Belotti 2015), second-order cone programming (SOCP) (Yamashita *et al*. 2018), matrix algebra computational tricks (Hinrichs *et al*. 2006), comparing the effect of kinship matrices on genetic gain, inbreeding and diversity (Gautason *et al*. 2023), including mate allocation optimization (Waldmann 2025) and various alternative formulations of the optimization problem to fit a range of different applications (Hallander and Waldmann 2009; Clark *et al*. 2013; Hamilton 2020; Hjortø *et al*. 2022; Fogg *et al*. 2024). Computational efficiency is of particular importance for the use of OCS in large-scale national and international breeding programs, where pedigrees can comprise millions of individuals, such as the Interbull international cattle genetic evaluation system (Weigel *et al*. 2001), the Breedbase platform used in plant breeding (Hershberger *et al*. 2021), and TreePlan in forest tree breeding (McRae *et al*. 2003), making single-solve optimization approaches preferable to methods that require repeated optimization in many scenarios.

The efficiency of OCS relies heavily on the accuracy of the estimated breeding values (EBVs) used to calculate the contributions of breeding candidates. Typically, EBVs are obtained as point estimates that are plugged directly into OCS, so that uncertainty in those estimates is not accounted for in the optimization and few studies have investigated this aspect. Fogg *et al*. (2024) introduced a robust OCS framework drawing on concepts from robust optimization, in which each candidate’s EBV uncertainty is represented by its standard error (SE), obtainable from frequentist inference or as a summary statistic from Bayesian methods. The robust OCS penalizes candidates whose high EBVs are accompanied by large SEs, preferring individuals with more reliable estimates even at some cost to expected gain; as the risk-aversion parameter approaches its maximum, this framework reduces to a minimax worst-case formulation that guards against the single most adverse realization of EBV uncertainty. Pocrnic *et al*. (2023) proposed a complementary probabilistic approach, applying OCS across posterior draws of EBVs from a Bayesian MCMC analysis of a simulated breeding population. Their results revealed considerable bias in the optimized contributions of standard OCS relative to the probabilistic approach, with the latter producing solutions closer to the true optimum, demonstrating that ignoring the uncertainty of EBVs in the selection process can lead to suboptimal decisions, even when the point estimates themselves are unbiased. However, the approach of Pocrnic *et al*. (2023) was evaluated on a small simulated pedigree, leaving open whether ignoring EBV uncertainty systematically compromises genetic progress in operational breeding populations (Bijma 2012), and whether a single-solve uncertainty-aware formulation, rather than repeated per-draw optimization, can achieve equivalent or superior protection at a fraction of the computational cost. We refer to the point-estimate approach throughout as MAP-OCS (maximum a posteriori OCS).

Both approaches above characterize EBV uncertainty marginally: robust OCS based on SE (Fogg *et al*. 2024) and per-draw posterior sampling (Pocrnic *et al*. 2023) treat each candi-date’s uncertainty independently, leaving unexploited a critical property of Bayesian MCMC genetic evaluation, i.e. the joint structure of the posterior (Sorensen and Gianola 2002). The full posterior *p*(**g** | **y**) captures not only the marginal variance of each individual’s breeding value but also the pairwise prediction error covariances (PEVs) arising from shared relatives, overlapping phenotypic information and genomic relationships (Henderson 1975). By sampling directly from this joint posterior at each MCMC iteration, the full PEV covariance structure propagates implicitly into the contribution optimization. Two candidates with positively correlated EBV uncertainty are jointly riskier to select together than two candidates with the same marginal SE whose uncertainties are independent or negatively correlated, a distinction that becomes especially consequential in populations with strong family structure or high genomic relatedness among elite candidates.

Exploiting this joint structure calls for a framework designed for decision-making under distributional uncertainty: portfolio optimization, in which the goal is to identify an allocation, consisting of a vector of parental contributions, that performs well not just under a single assumed scenario but robustly across the full distribution of possible outcomes (Markowitz 1952). In financial portfolio theory, the standard tool for quantifying and minimizing downside risk is the *Conditional Value at Risk* (CVaR), also known as Expected Shortfall (Rockafellar and Uryasev 2000, 2002). CVaR at confidence level *α* measures the expected loss in the worst (1 − *α*) fraction of scenarios. In the OCS context, this corresponds to the expected genetic gain when breeding values fall in their least favorable tail across MCMC posterior draws. Crucially, CVaR is a *coherent* risk measure (Artzner *et al*. 1999), meaning it satisfies subadditivity. Hence, diversification across individuals can only reduce, never increase, tail risk. This makes CVaR particularly well suited to OCS, where contribution diversification is already the central mechanism for managing inbreeding. CVaR-OCS directly addresses the limitations of both prior approaches. Unlike the SE-based robust OCS of Fogg *et al*. (2024), which treats each candidate’s uncertainty independently and marginally, CVaR-OCS propagates the full joint posterior uncertainty into the optimization through the MCMC scenario set, and subsumes the Fogg *et al*. (2024) framework as a limiting case when the confidence level *α* approaches one, and in contrast to the per-draw approach of Pocrnic *et al*. (2023), CVaR-OCS requires solving only a single quadratic program rather than one per posterior sample, making it computationally practical for operational breeding programs with large pedigrees.

The aim of our study is threefold. First, we introduce CVaR-OCS, a novel formulation of optimum contribution selection that directly incorporates the joint posterior distribution of EBVs into the optimization objective via Conditional Value at Risk (CVaR). CVaR-OCS solves a single quadratic program that simultaneously maximizes expected genetic gain and minimizes tail risk arising from EBV uncertainty, providing a principled and computationally efficient alternative to point-estimate OCS that requires no repeated optimization across MCMC samples. Second, we use Monte Carlo sampling across MCMC iterations (MCMC-OCS) as a diagnostic tool to characterize how EBV uncertainty propagates into contribution decisions, quantifying the instability of MAP-OCS selections through overlap metrics, selection frequency analysis, and individual robustness scores. These diagnostics motivate CVaR-OCS and provide complementary insight into the sources of selection instability. Third, we evaluate CVaR-OCS and its MCMC-OCS diagnostics on a multi-generation genomic selection simulation using the QTLMAS 2010 benchmark dataset (Szydłowski and Paczyńska 2011) with known true breeding values, enabling direct comparison to an oracle solution and assessment of decision quality under controlled EBV uncertaintyand on an operational Norway spruce (*Picea abies*) forest tree breeding pedigree.

## Materials and methods

### Optimum contribution selection

Let *n* denote the number of individuals in the recruitment or candidate population and let **Σ** ∈ ℝ^*n*×*n*^ be the matrix of pairwise genetic relationships among candidates, inferred from pedigree information, marker data, or a combination of both (see Case studies below for the specific matrix used in each analysis). For optimum contribution selection (OCS), the optimization problem can be formulated as (Pong-Wong and Woolliams 2007; Ahlinder *et al*. 2013; Mullin and Belotti 2015; Yamashita *et al*. 2018):

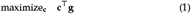

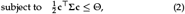

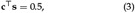

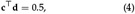

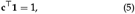

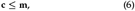

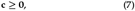

where **c** ∈ ℝ^*n*^ is the vector of individual contributions (the decision variable), **g** ∈ ℝ^*n*^ is the vector of estimated breeding values (EBVs) of the selection candidates, Θ is the upper bound on group coancestry of the selected cohort, **m** is a vector of upper bounds on the contribution of any single individual arising from practical limitations, and **s, d** are indicator vectors for sires and dams, respectively. Note that the gender constraints (3)–(4) are included for dioecious species and removed when the species is monoicous, in which case the total-contribution constraint (5) ensures **c**^⊤^ **1** = 1. The objective function (1) is linear in **c** with a quadratic constraint on average relatedness among the selected candidates, so the problem (1)–(7) is a quadratic programme (QP). Here, we used the COSMO solver (Garstka *et al*. 2019) available in the JuMP modelling language for Julia (Dunning *et al*. 2017). This solver handles convex optimization problems with quadratic objective functions and conic constraints using an operator splitting algorithm based on ADMM (Boyd *et al*. 2010). Please consult Supplementary information for further details on the tested optimization methods and computational performance on the analyzed pedigrees.

### Bayesian mixed model and MCMC inference

Following Sorensen and Gianola (2002), we used the following multi-trait mixed effects model to infer breeding values:

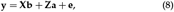

where **y** is the vector containing each of *m* traits represented sequentially for *n* individuals in a single column vector of size *nm* × 1; **X** and **Z** are the fixed and random-effect design matrices of sizes *nm* × *pm* and *nm* × *nm*, respectively, associating predictors with the corresponding phenotypic records for *p* levels of fixed-effect predictors. Random polygenic and residual effects are assumed to follow, for two traits, one categorical and one continuous:

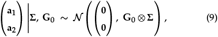

where **a**_1_ and **a**_2_ are the vectors of breeding values (polygenic scores) for the continuous trait (subscript 1) and the categorical trait (subscript 2), respectively. In addition, **Σ** is the *n* × *n* genetic relationship matrix of the candidates (pedigree **A**, genomic **G**, or blended **H** as specified per case study) and **G**_0_ is the 2 × 2 genetic covariance matrix between traits, defined as

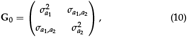

where 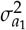 and 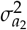 are the additive genetic variances for traits 1 and 2, and 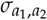 is their additive genetic covariance. The joint phenotypic covariance is Cov(**y**) = **Z**(**G**_0_ ⊗ **Σ**)**Z**^⊤^ + **R**, where **R** is the residual covariance matrix defined analogously to **G**_0_ but with error (co)variances. For the categorical trait (trait 2), a threshold model is assumed on the underlying liability scale (Sorensen and Gianola 2002), where the observed categorical response is determined by an underlying continuous liability exceeding threshold values. Prior distributions were assigned following the default structure of JWAS (Cheng *et al*. 2018). Fixed effects intercept, trial (site), and plot nested within trial received flat, improper priors *p*(***β***) ∝ 1, placing no constraint on location parameters. The plot-within-trial term was treated as an i.i.d. random effect with a scaled inverse chi-squared prior on its variance, with degrees of freedom *ν* = 4.0 (JWAS default) and scale derived from the phenotypic variance of the corresponding trait.

For the multivariate additive genetic component under GBLUP, the genetic covariance matrix **G**_0_ was assigned an inverse-Wishart prior,

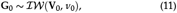

with degrees of freedom *ν*_0_ = 4.0 and scale matrix **V**_0_ derived from the phenotypic (co)variances of the response traits, consistent with the JWAS default when no explicit prior scale is supplied (Cheng *et al*. 2018). The residual covariance matrix **R** was assigned an analogous inverse-Wishart prior with the same default parameterisation. No informative prior knowledge was encoded for any variance component.

For the H-matrix model, threshold formulation was assumed for the categorical trait Lev17 (tree survival, binary) on the underlying liability scale (Sorensen and Gianola 2002). Observed binary responses were recoded to category labels {1, 2} as required by JWAS, and liability values were imputed at each Gibbs iteration by data augmentation (Cheng *et al*. 2018). The liability-scale residual variance for Lev17 was fixed at 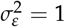 for model identifiability, which is the standard constraint in threshold models and removes one degree of freedom from the residual variance prior for that trait; the threshold separating the two liability categories was fixed at zero. The three continuous traits (Hjd17, Htv17, Sprant17) retained freely estimated residual variances under the same inverse-Wishart prior structure as the G-matrix model.

After inferring breeding values **a**_1_ and **a**_2_, a joint selection index matrix **I** of size *n* × *l* was defined for all members of the *n*- individual pedigree across *l* MCMC iterations by weighting the breeding values of the traits. For simplicity, all included traits were given equal weights. The multi-trait mixed effects model was implemented in the JWAS package (Cheng *et al*. 2018) in Julia using default settings, with the total chain length of 52,000 iterations, a burn-in of 2,000 iterations, and thinning by a factor of 10, yielding *l* = 5,000 post-burn-in posterior samples for all subsequent OCS analyses. Chain convergence was assessed by visual inspection of trace plots and by computing the effective sample size (ESS) for all heritability and genetic variance parameters using the MCMCDiagnosticTools.jl package; convergence statistics are reported in Supplementary materials.

### MCMC-based uncertainty analysis

Before introducing the CVaR-OCS optimization framework, we first characterize how EBV uncertainty propagates into contribution decisions by solving OCS separately for each of the *l* MCMC posterior draws. This repeated sampling procedure (referred to as MCMC-OCS throughout) serves as a diagnostic tool. The resulting distribution of contribution vectors {**c**^(1)^, …, **c**^(*l*)^ } reveals which MAP-OCS selections are stable across the posterior and which are sensitive to EBV uncertainty, directly motivating the single-solve CVaR-OCS formulation described in Section Conditional Value at Risk. Formally, for each MCMC iteration *j* we solve:

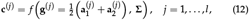

where *f* (·) is the OCS function call and 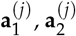 are the EBVs of the selection candidates for the continuous and categorical traits in iteration *j*. For the general m-trait case the index weights **w** replace the 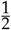 scalar. Using this procedure, we acquire a matrix of contributions **C** = [**c**^(1)^ **c**^(2)^ · · · **c**^(*l*)^ ].

#### MAP-OCS reference solution

The reference optimum contribution vector **c**^MAP^ was computed using maximum a posteriori (MAP) breeding values:

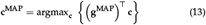

subject to constraints (2)–(7), where **g**^MAP^ is the vector of MAP breeding values.

#### Overlap and agreement metrics

For each MCMC iteration *j*, the top 100 contributors were identified and the overlap with the MAP-OCS solution was computed as

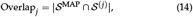

where 𝒮^MAP^ denotes the set of top 100 MAP-OCS contributors and 𝒮 ^(*j*)^ denotes the top 100 contributors from iteration *j*. Jaccard similarity was computed as

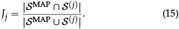

#### EBV–selection frequency analysis

Selection frequency for each individual across all MCMC iterations was calculated as

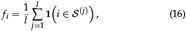

where **1**(*i* ∈ 𝒮 ^(*j*)^) is an indicator function equal to 1 if individual *i* is in the top 100 contributors in iteration *j*. The relationship between MAP EBV and MCMC-OCS selection frequency was analysed using Gaussian process (GP) regression (Rasmussen and Williams 2006) with a Matérn 3/2 covariance function, with all hyperparameters estimated via maximum likelihood using the Nelder–Mead algorithm. The GP model was fitted using AbstractGPs.jl (van der Wilk *et al*. 2020) and Optim.jl (Mogensen and Riseth 2018) in Julia 1.11.

### Stochastic programming interpretation of MCMC-OCS

The OCS problem (1)–(7) treats the breeding value vector **g** as a fixed, known quantity. In reality **g** is a random vector whose uncertainty is fully characterised by the posterior distribution *p*(**g** | **y**) obtained from the Bayesian MCMC analysis described above. Ignoring this uncertainty can lead to suboptimal and fragile selection decisions (Fogg *et al*. 2024; Pocrnic *et al*. 2023). A natural framework for making decisions under such uncertainty is *stochastic programming* (Birge and Louveaux 2011; Shapiro *et al*. 2009), in which uncertain parameters are represented by a probability distribution and the optimisation objective is defined as an expectation (or a risk-adjusted expectation) over that distribution.

#### Scenario-based formulation

Let Ω denote the probability space of breeding values and let ***ξ***(*ω*) = **g**(*ω*) be the random breeding-value vector for outcome *ω* ∈ Ω. The general mean-risk stochastic OCS problem is

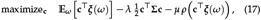

subject to the feasibility constraints (3)–(7), where *λ* > 0 is the inbreeding-penalty weight replacing the hard constraint (2) with a Lagrangian penalty (these two formulations trace the same efficient frontier as *λ* varies), *µ* ≥ 0 is a risk-aversion weight, and *ρ*(·) is a chosen risk measure applied to the distribution of realised genetic gain **c**^⊤^ ***ξ***(*ω*).

The MCMC sampler produces *l* approximately independent draws {**g**^(1)^, **g**^(2)^, …, **g**^(*l*)^ } from the posterior *p*(**g** | **y**). Each draw constitutes a *scenario*, and the Sample Average Approximation (SAA) (Shapiro *et al*. 2009; Klein Haneveld *et al*. 2015) replaces the expectation and the risk measure by their empirical counterparts over the *l* scenarios

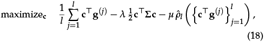

subject to (3)–(7), where 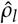 is the empirical risk measure over the *l* scenarios. When *µ* = 0 the objective reduces to 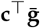, where 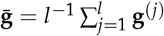 is the posterior mean EBV, and the problem is reduced to a standard QP equivalent to MAP-OCS evaluated at the posterior mean.

#### Relationship to the MCMC-OCS procedure

The MCMC-OCS procedure implemented in the present study (following Pocrnic *et al*. 2023) solves *l* separate OCS problems, one per MCMC draw (12), and analyses the resulting distribution of contribution vectors {**c**^(1)^, …, **c**^(*l*)^ } . This differs fundamentally from the joint SAA problem (18). In our procedure the contribution vector **c** is reoptimised for each scenario rather than held fixed across all scenarios, so EBV uncertainty appears in the output distribution of **c** but not in the optimisation itself. The joint SAA (18), described here for theoretical completeness, identifies a single robust **c** that explicitly trades off expected gain, inbreeding and estimation risk simultaneously. This formulation is the basis for the CVaR-OCS extension introduced below.

#### Connection to Markowitz mean–variance theory

Formulation (18) has an instructive connection to Markowitz mean–variance portfolio theory (Markowitz 1952, 1959). In the Markowitz problem, the variance constraint is relaxed into the objective via its Lagrange multiplier, yielding the penalised form

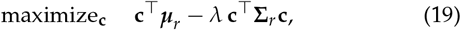

where ***µ***_*r*_ is the vector of expected returns and **Σ**_*r*_ is their covariance matrix. The OCS objective (18) (with *µ* = 0) is structurally identical, with the posterior mean 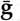 playing the role of expected returns and **Σ** playing the role of the return covariance matrix. The dual formulation, which maximises the expected genetic gain for a given inbreeding budget, traces the same efficient frontier. In addition, the Lagrange multiplier *λ* is the risk of inbreeding, analogous to the market price of risk in finance. This connection motivates the portfolio-level vulnerability metrics described in the Supplementary Material (Section S2.2), which are conceptually equivalent to portfolio diversification measures (Markowitz 1952).

#### Value at Risk

When breeding values are uncertain, the genetic gain *Z*(**c, *ξ***) = **c**^⊤^ ***ξ*** is itself a random variable. A natural summary of the downside of this distribution is the *Value at Risk* (VaR) at confidence level *α* ∈ (0, 1), defined as the *α*-quantile of the gain distribution

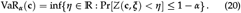

Intuitively, VaR_*α*_ (**c**) is the level of genetic gain that will be exceeded with probability *α*. For example, with *α* = 0.95, the VaR is the genetic gain that will *not* be achieved only 5% of the time across the posterior distribution of breeding values. In the OCS context, a low VaR_*α*_ for a given contribution vector **c** signals that the selection decision is sensitive to EBV uncertainty. Even moderate perturbations of the breeding values can substantially reduce genetic gain. In the SAA setting, VaR_*α*_ is estimated by the empirical *α*-quantile of the sample 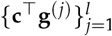.

Despite its intuitive appeal, VaR has well-known limitations as a risk measure. It conveys no information about the severity of outcomes beyond the threshold. Furthermore, VaR is not in general a coherent risk measure (Artzner *et al*. 1999) since it lacks the subadditivity property, so combining two contribution vectors can increase rather than decrease VaR, which is an undesirable feature for a diversification-based framework such as OCS.

#### Conditional Value at Risk

These limitations are addressed by the *Conditional Value at Risk* (CVaR), also known as Expected Shortfall (Rockafellar and Uryasev 2000, 2002). CVaR at level *α* is defined as the expected genetic gain conditional on being in the worst (1 − *α*) fraction of the posterior distribution:

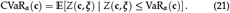

CVaR is a coherent risk measure (Artzner *et al*. 1999) and is therefore well suited to the OCS diversification objective. Maximising CVaR_*α*_ (**c**) is equivalent to maximising the expected genetic gain in the worst (1 − *α*) scenarios, providing explicit protection against cases where the true breeding values are substantially lower than their posterior means.

A key computational advantage of CVaR is the Rockafellar and Uryasev (2000) auxiliary-variable representation

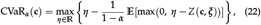

where *η* equals VaR_*α*_ (**c**) at the optimum. Substituting the SAA estimator for the expectation and reinstating the hard inbreeding constraint gives the following tractable QP for CVaR-aware OCS

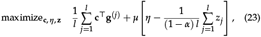

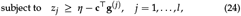

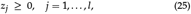

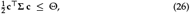

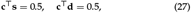

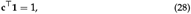

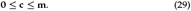

Here *z*_*j*_ = max(0, *η* − **c**^⊤^ **g**^(*j*)^) is the shortfall of genetic gain in scenario *j* below the VaR threshold *η*, and *µ* ≥ 0 controls the weight given to tail-risk protection. Note that the inbreeding penalty of (17) has been replaced by the hard constraint (26) with threshold Θ, consistent with the original OCS formulation (2). Thus *µ* is therefore the sole risk-aversion parameter. As in (3)– (4), the gender constraints (27) are included only for dioecious species.

Problem (23)–(29) is a QP in (**c**, *η*, **z**), quadratic in **c** through (26), and linear in *z*_*j*_ and *η*, and can be solved directly with the COSMO solver (Garstka *et al*. 2019) in JuMP (Dunning *et al*. 2017). The objective has a clear two-component interpretation: the first term maximises expected genetic gain across all posterior scenarios (analogous to the expected-return term in Markowitz), while the CVaR term penalises poor outcomes in the tail of the breeding-value distribution, capturing candidate-specific estimation risk. As *µ* → 0 the CVaR term vanishes and the problem reduces to MAP-OCS at the posterior mean, while as *α* → 1, CVaR converges to the worst-case scenario, approaching the minimax robust OCS of Fogg *et al*. (2024). The parameter *µ* therefore spans a risk–return continuum that generalises both MAP-OCS and the robust OCS of Fogg *et al*. (2024) as limiting cases.

### CVaR-OCS parameter selection

The CVaR-OCS formulation (23)–(29) involves two parameters that jointly determine the operating point on the risk–return efficiency frontier, the confidence level *α* ∈ (0, 1) and the risk-aversion weight *µ* ≥ 0.

The confidence level *α* defines which tail of the gain distribution is protected. At *α* = 0.90, CVaR-OCS optimises the expected genetic gain in the worst 10% of MCMC scenarios, and at *α* = 0.99 it focuses exclusively on the worst 1%. Higher *α* therefore provides stronger tail protection but concentrates the optimisation objective on an increasingly small and extreme subset of posterior draws, which can reduce stability when the number of MCMC iterations *l* is modest. The risk-aversion weight *µ* governs the relative importance of tail-risk protection versus expected genetic gain. At *µ* = 0 the CVaR term vanishes and the problem reduces to MAP-OCS evaluated at the posterior mean (equation 23), while increasing *µ* progressively shifts the solution toward the minimax worst-case limit (Fogg *et al*. 2024).

To select a practically useful operating point, we evaluated all combinations of *α* ∈ {0.90, 0.95, 0.99} and *µ* ∈ {0.50, 0.80, 1.00, 1.20, 1.50, 1.80, 2.00, 5.00, 10.00} for each dataset. For each (*α, µ*) combination, we recorded the percentage genetic gain loss and the percentage 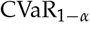 improvement relative to the MAP-OCS baseline, yielding a gain–tail-risk efficiency frontier for each *α* level.

The recommended operating point on each frontier was identified using a geometric perpendicular-distance criterion (see also Satopaa *et al*. 2011). Each frontier was normalised to [0, 1] in both dimensions and the point of maximum perpendicular distance from the line connecting the two endpoints was taken as the elbow, i.e. the point beyond which additional tail-gain improvement comes at disproportionately high genetic gain cost. As a confirmation, we computed the marginal efficiency ratio between successive *µ* values, defined as the ratio of 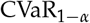 improvement to loss of genetic gain. A sharp drop in this ratio confirms that the elbow marks a genuine change in the efficiency regime rather than a gradual transition. The selected (*α, µ*) combination was used for all downstream comparisons within each dataset.

### MCMC-based robustness scores and contribution distribution

To characterize which MAP-OCS selections are most vulnerable to EBV uncertainty, we calculated individual robustness scores by quantifying the expected loss of genetic gain when each candidate is excluded from the selected cohort across MCMC iterations. We also expressed gain as a percentage relative to the mean baseline, providing a more stable normalized metric. High-risk individuals were then identified using a dynamic bottom-quartile threshold applied to MAP-OCS selections. To characterize population-level genetic gain concentration, we computed four complementary contribution distribution metrics: mean marginal gain, maximum marginal gain, top-10% concentration ratio, and the Gini coefficient. Full definitions, equations, and computational details are provided in the Supplementary Material.

### Simulated animal breeding dataset

The simulated dataset from the 14th QTL-MAS workshop (Szydłowski and Paczyńska 2011) was used to validate CVaR-OCS in an animal breeding context with explicit sex structure and known true breeding values (TBVs). The pedigree comprised *n* = 3,226 individuals across five generations, founded by 5 sires and 15 dams. Two correlated traits, a quantitative trait and a binary trait, were simulated under a model including additive, epistatic, and parent-of-origin effects.

Phenotypic records were available for *n*_pheno_ = 2,326 individuals from the first four generations, while the *n*_cand_ = 900 individuals of the fifth generation were un-phenotyped and served as the candidate population for OCS, mimicking a standard genomic selection scenario in which selection candidates lack own performance records. Genotypic data comprised *p* = 9,723 SNPs on five chromosomes after quality control (MAF > 0.05, missingness < 10%). Sex was recorded for all individuals and included as a fixed effect in the Bayesian model.

GEBVs and their full posterior distributions were estimated for all *n* = 3,226 individuals using a bivariate Bayesian model in JWAS (Cheng *et al*. 2018), with the genomic relationship matrix **G** constructed following VanRaden (2008) for use in the OCS coancestry constraint. The posterior MCMC samples of GEBVs for the *n*_cand_ = 900 fifth-generation candidates were subsequently used as input to both MAP-OCS (using the posterior mean as a point estimate) and MCMC-OCS. Because the true additive breeding values of all individuals are known by construction in this dataset, the selections recommended by each method could be evaluated against the TBV-optimal solution, which provides a direct assessment of decision quality under EBV uncertainty.

### Norway spruce case study

Two field trials of Norway spruce, with Skogforsk IDs S23F8820476 located outside Vindeln, Västerbotten County (64.3^°^Latitude, 19.68^°^Longitude) and S23F8820477, Hädanberg, Västernorrland County (63.58^°^Latitude, 18.20^°^Longitude), were utilized for illustrating CVaR-OCS and MCMC-OCS on forest tree breeding field trials (Table 1). In both trials, approximately 6000 offspring from 150 families were planted in 1988 in single tree plot design for evaluating the performance of parent trees in the Hissjö seed orchard. The following phenotypic traits were scored: total tree height at year 7 and 17, early height growth at year 17, number of spike knots at year 17 and tree vitality at year 7. Of those traits, tree height, height growth and number of ramicorns at age 17 were used in the selection index with equal weight (negative weight for ramicorn number). In total, 1218 offspring (568 and 650 in S23F8820476 and S23F8820477 respectively) where genotyped using exome capture technique resulting in 300 714 SNPs prior to filtration. For further details of the Norway spruce data, please consult Chen *et al*. (2018). The G matrix was calculated using the rrBLUP r package (Endelman 2011), with default settings. To combine genotyped and ungenotyped trees, a joint **H** matrix was calculated using the AGHmatrix r package (Amadeu *et al*. 2023), where **G** was first tuned towards **A** following Christensen *et al*. (2012) by matching mean diagonal and off-diagonal elements (tuning coefficients *α* = 1.025, *β* = 0.029). The blending was performed using the Hmatrix function, with *τ* = 1 and *ω* = 1 (default) (Legarra *et al*. 2009). To ensure that **H** used in subsequent analyses was positive definite, we applied the cor_nearPD function from the Bigsimr.jl package. Multi-trait analysis to obtain posterior estimates of breeding values (BV) were performed using the JWAS Julia package (Cheng *et al*. 2018) with a total chain length of 52,000 iterations, a burn-in of 2,000 iterations, and thinning by a factor of 10, yielding *l* = 5,000 post-burn-in posterior samples. The predictors used in the model were a site factor and a nested plot within site term.

**Table 1.** Description of included datasets. Average coancestry (Avg. coanc.) of the candidate population is calculated via the pedigree.

| Study | Species | Pedigree | Candidates | Gen. | Parents | Families | Avg. coanc. |
| --- | --- | --- | --- | --- | --- | --- | --- |
| Szydlowski and Paczyńska (2011) | simulated animals | 3,226 | 900 | 4 | 59 | 30 | 0.054 |
| Chen *et al.* (2018) | *P. abies* | 5,597 | 5,525 | 2 | 72 | 146 | 0.016 |
| Chen *et al.* (2018) | *P. abies* | 1,273 | 1,218 | 2 | 55 | 132 | 0.014 |

## Results

### Validation on a simulated dataset with known true breeding values

To evaluate the new proposed framework CVaR-OCS under controlled conditions where true breeding values (TBVs) are known, we analysed the QTL-MAS 2010 workshop dataset (Szydłowski and Paczyńska 2011), which provides a realistic multi-generation genomic selection scenario with a fully genotyped pedigree. The final generation of *n* = 900 unphenotyped selection candidates (433 sires, 467 dams) served as the target population for OCS, with a primary coancestry constraint of Θ = 0.03. Posterior distributions were obtained from a two-trait Bayesian GBLUP model fitted in JWAS with *l* = 5000 post-burnin MCMC samples. An oracle solution, MAP-OCS(TBV), computed using known TBVs in place of EBVs, served as the theoretical optimum against which both EBV-based methods were evaluated. The bivariate JWAS model estimated moderate heritabilities for both traits (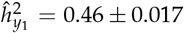 for the continuous trait; 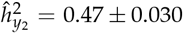 for the binary trait) and a positive inferred genetic correlation of 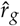 = 0.69, indicating that the two traits share substantial genetic architecture and that a combined selection index with equal weights captures genetic merit across both simultaneously.

#### CVaR-OCS parameterisation and elbow selection

To identify the operating point that appropriately penalises EBV uncertainty without sacrificing excessive genetic gain, we evaluated all combinations of *α* ∈ {0.90, 0.95, 0.99} and *µ* ∈ {0.5, 0.8, 1.0, 1.2, 1.5, 1.8, 2.0, 5.0, 10.0} across three coancestry constraints Θ ∈ {0.02, 0.03, 0.05}, with results at Θ = 0.03 presented in the main text and the remaining constraints in Supplementary Table S7. The efficiency frontier (Figure 1a), expressed as percentage genetic gain loss and CVaR_95_ improvement relative to the MAP-OCS in-sample baseline, reveals a consistent concave relationship. Early increments in *µ* yield disproportionate tail-gain improvements, with returns diminishing beyond a species-specific elbow.

**Figure 1.**
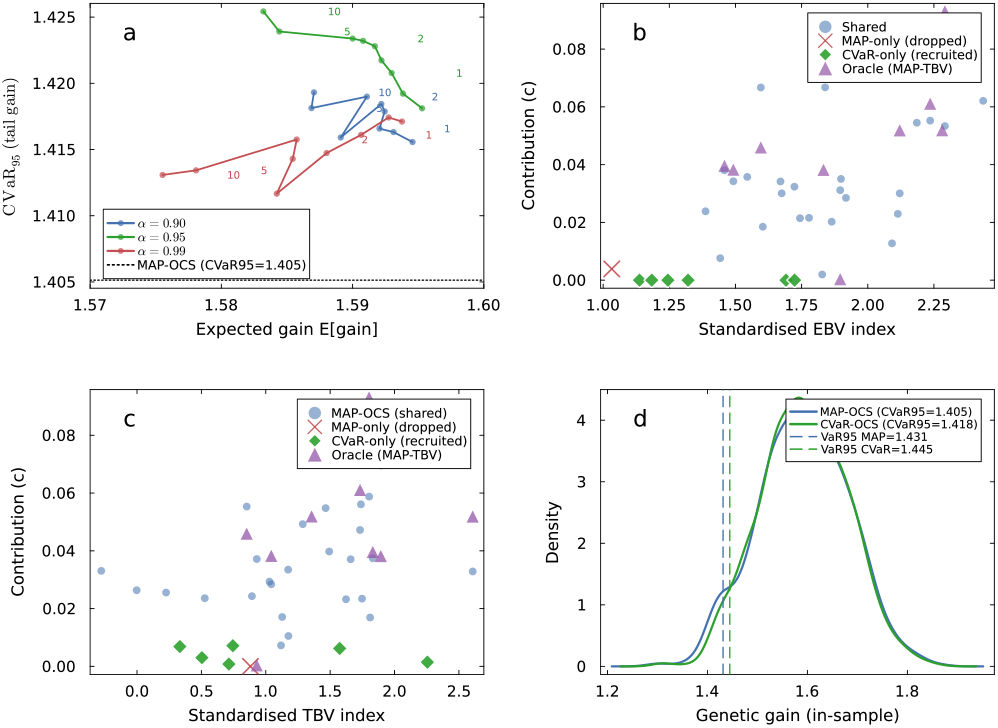
CVaR-OCS validation on the QTLMAS 2010 simulated dataset (*n* = 900 candidates; Θ = 0.03). **a** Gain–tail-risk efficiency frontier across three confidence levels (*α* 0.90, 0.95, 0.99 ) and CVaR weights (*µ* 0.50, …, 10.00 ), expressed as percentage genetic gain loss and CVaR_95_ improvement relative to the MAP-OCS in-sample baseline. Open diamonds mark the elbow point for each *α* level, identified by the geometric perpendicular-distance criterion (*CVaR-OCS parameter selection* in Materials and Methods), beyond which additional CVaR_95_ improvement comes at disproportionate genetic gain cost. **b** Standardised EBV index vs parental contribution for MAP-OCS and the recommended CVaR-OCS solution (*α* = 0.95, *µ* = 1.50), coloured by selection group: individuals shared between MAP-OCS and CVaR-OCS (blue circles), the single MAP-OCS selection dropped by CVaR-OCS (red cross), and individuals newly recruited by CVaR-OCS (green diamonds). The oracle solution based on true breeding values (MAPOCS(TBV); purple triangles) is shown for reference. **c** Standardised TBV index vs parental contribution for the same three solutions as in panel **b**. CVaR-OCS recruits individuals whose true breeding values more closely align with the oracle solution compared to MAP-OCS (EBV-based), demonstrating improved robustness to EBV uncertainty when true genetic merit is considered. **d** Kernel density of genetic gain realised by MAP-OCS (blue) and CVaR-OCS (green) evaluated across all *l* = 5000 MCMC EBV scenarios. Dashed vertical lines mark the 5th-percentile gain (VaR_95_) for each solution; CVaR_95_ values are given in the legend. CVaR-OCS improves tail-gain security relative to MAP-OCS while maintaining similar mean expected gain, consistent with the Norway spruce results (Figure 4).

Elbow points were identified using the geometric perpendicular-distance criterion (*CVaR-OCS parameter selection* in Materials and Methods): *α* = 0.90 at *µ* = 1.75 (CVaR_95_ +0.77%, gain loss +0.13%), *α* = 0.95 at *µ* = 1.50 (CVaR_95_ +1.26%, gain loss − 0.04%), and *α* = 0.99 at *µ* = 1.50 (CVaR_95_ +0.46%, gain loss +0.43%; Table 2). The recommended operating point is *α* = 0.95, *µ* = 1.50, which achieves the highest CVaR_95_ improvement (+1.26%) at negligible gain cost (− 0.04% vs. the MAP-OCS in-sample baseline; Figure 1a). *α* = 0.99 produced consistently lower CVaR_95_ improvements than *α* ∈ {0.90, 0.95} at all *µ* values, reflecting the noisier tail estimate at this confidence level (only 40 tail draws at *l* = 5000), and confirms that extreme confidence levels are counterproductive when *l* is moderate.

**Table 2.** **CVaR-OCS results for the QTL-MAS 2010 simulation study** at the recommended operating point (Θ = 0.03; *α*^∗^ = 0.90, *µ*^∗^ = 1.8; *n* = 900 candidates; *l* = 5,000 MCMC scenarios). *E*[gain]: in-sample expected genetic gain^a^; Realised gain: **c**^⊤^ **t** evaluated against true breeding values (TBVs); CVaR_95_: expected gain in the worst 5% of MCMC scenarios; *n*_sel_: number of selected parents (sires S, dams D); Jaccard: set overlap with oracle. Results across all three coancestry constraints (Θ ∈ { 0.02, 0.03, 0.05 }) are provided in Supplementary Table S7.

| Metric | Solution | Value | $\Delta(\%)$ |
| --- | --- | --- | --- |
| $E[\text{gain}]$ | Oracle | 1.725 | — |
|  | MAP-OCS(EBV) | 1.595 | — |
|  | CVaR-OCS | 1.596 | +0.1 |
| Realised gain | MAP-OCS(EBV) | 1.342 | — |
|  | CVaR-OCS | 1.325 | −1.2 |
| $\text{CVaR}_{95}$ | MAP-OCS(EBV) | 1.400 | — |
|  | CVaR-OCS | 1.437 | +2.7 |
| $\text{VaR}_{95}$ | MAP-OCS(EBV) | 1.440 | — |
|  | CVaR-OCS | 1.469 | +2.1 |
| $n_{\text{sel}}$ (S, D) | Oracle | 31 (15, 16) | — |
|  | MAP-OCS(EBV) | 29 (17, 12) | — |
|  | CVaR-OCS | 35 (20, 15) | +20.7 |
| Jaccard vs oracle | MAP-OCS(EBV) | 0.176 | — |
|  | CVaR-OCS | 0.196 | +11.4 |
| Shared with oracle | MAP-OCS(EBV) | 9 | — |
|  | CVaR-OCS | 11 | +22.2 |

<sup>a</sup> MAP-OCS(EBV) $E[\text{gain}]$ is optimised against the posterior-mean EBV vector and overestimates realised gain by 18.8% at $\Theta = 0.03$ , due to EBV uncertainty. CVaR-OCS $E[\text{gain}]$ is jointly optimised over all $l$ scenarios and carries no such bias.

#### Tail-risk protection and contribution structure

Having selected *α* = 0.95, *µ* = 1.50 as the recommended operating point, we examined what CVaR-OCS actually delivers relative to MAP-OCS in terms of tail-gain security and the structure of parental contributions. At *α* = 0.95, *µ* = 1.50, CVaR-OCS improved CVaR_95_ by 0.91% relative to MAP-OCS(EBV) (1.4051 → 1.4179) and VaR_95_ by 0.94% (1.4312 → 1.4447) when both solutions were evaluated across all *l* = 5000 MCMC scenarios (Figure 1d; Table 2). The gain distribution variance was reduced by 4.8% under CVaR-OCS (F-test: *F* = 1.10, *p* = 0.081; Cohen’s *d* = 0.015), with no significant change in mean gain (t-test: *p* = 0.76), indicating that CVaR-OCS compresses the lower tail of the distribution without affecting the centre.

CVaR-OCS selected 35 parents compared to 28 under MAP-OCS (+25% broader selection base), distributing contributions more equitably (Gini: 0.389 vs. 0.326). Of the 35 CVaR-OCS selections, 27 were shared with MAP-OCS, one MAP-OCS individual was dropped, and 8 new individuals were recruited (Figure 1b). The single dropped individual illustrates the core mechanism: CVaR-OCS identified one MAP-selected animal whose contribution to genetic gain was not robust across MCMC scenarios and replaced it with eight moderate-EBV individuals whose portfolio-level diversification reduced tail sensitivity.

#### Estimation bias in MAP-OCS

A pronounced discrepancy was observed between the genetic gain that MAP-OCS(EBV) claims when optimised against posterior-mean EBVs and its realised gain when evaluated against the full MCMC scenario distribution. At Θ = 0.03, MAP-OCS(EBV) reported an optimised gain of 1.91 (vs. posterior-mean EBVs), but realised only 1.59 in-sample across MCMC scenarios, which is an overestimation of 16.7%. This bias arises because MAP-OCS treats the posterior mean as a fixed, known quantity and selects a optimal population without EBV uncertainty. CVaR-OCS is free of this bias by construction, as it is jointly optimised over all *l* scenarios. Its in-sample expected gain (1.592 at *α* = 0.95, *µ* = 1.50) is directly comparable to the evaluation-based estimate without systematic inflation.

#### Oracle comparison

The oracle MAP-OCS(TBV) selected 31 individuals spanning a wider EBV range than either EBV-based method (Figure 1b,c), reflecting the oracle’s knowledge that several high-EBV individuals carry inflated posterior-mean estimates. Jaccard similarity between MAP-OCS(EBV) and the oracle was 0.180 (*n* = 9 shared individuals) at Θ = 0.03. CVaR-OCS improved oracle overlap marginally to 0.179 (*n* = 10 shared; +11% more oracle-optimal individuals recovered), despite having no access to true values.

### Application to an operational Norway spruce breeding population

#### Genetic parameter estimates

Heritability estimates varied markedly across relationship matrices and traits in Norway spruce (Supplementary Table S3). MCMC convergence was confirmed for all models, with chains sampling their respective stationary distributions as shown in the trace plots and posterior distributions (Supplementary Figures S1–S2 for the G-matrix model and S3–S4 for the H-matrix model). Genomic (**G**-matrix) heritabilities for the two primary height traits entering the selection index were nearly half the corresponding blended (**H**- matrix) estimates: 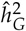 = 0.251 ± 0.048 vs. 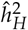 = 0.442 ± 0.045 for Htv17, and 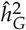 = 0.221 ± 0.056 vs. 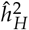 = 0.442 ± 0.040 for Hjd17. The third index component, ramicorn number at age 17 (Sprant17), showed very low genomic heritability (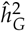 = 0.094 ± 0.040), indicating that uncertainty is compounded across all three index traits.

Posterior distributions of EBVs revealed substantial ranking uncertainty within families in Norway spruce (Fig. 2). In the high-performing family, ranking standard deviation reached 7.4, with the top-ranked individual having only 48% probability of ranking in the top 10% of its family. These overlapping posterior distributions illustrate the core challenge motivating our uncertainty-aware framework.

**Figure 2.**
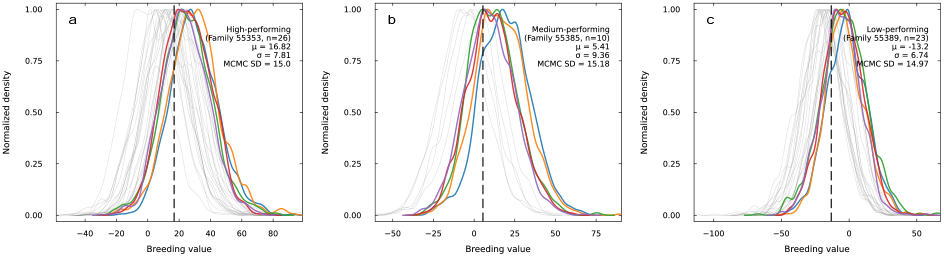
Posterior uncertainty in estimated breeding values reveals ranking ambiguity within Norway spruce families. Kernel density plots show the posterior distributions of selection index breeding values (Hjd17 + Htv17 −Sprant17) for individuals within three representative families: **a** high-performing (Family 55353, *n* = 26), **b** medium-performing (Family 55385, *n* = 10), and **c** low-performing (Family 55389, *n* = 23). Each thin curve represents the posterior distribution for one individual based on 5,000 MCMC samples. Thick curves highlight the top 5 individuals within each family ranked by posterior mean. Dashed vertical lines indicate family means.

#### MCMC-OCS characterizes the impact of EBV uncertainty on contributions

To characterize how EBV uncertainty propagates into OCS decisions, we solved OCS for each of the *l* MCMC posterior draws and compared the resulting distribution of contribution vectors to the MAP-OCS reference solution. This diagnostic procedure reveals the degree to which MAP-OCS selections are stable across the posterior distribution of breeding values and, by extension, which individuals are candidates for the risk-aware revision implemented by CVaR-OCS.

For the Norway spruce **G** population (*n* = 1,218), MCMC-OCS revealed considerable uncertainty in contributions, with a mean overlap of only 26.6 individuals (range: 10–46) and mean Jaccard similarity of 15.4% relative to MAP-OCS (Supplementary Table S8; Fig. 3a-d). Nearly all individuals were selected in at least some MCMC iterations, yet not a single one was selected in more than half of them (highest at 35.7% selection frequency, Fig. 3b). Despite this instability in individual allocations, a strong rank-order relationship between EBV and selection frequency was preserved (Spearman *ρ* = 0.932), with top-quartile EBV individuals selected 22.9× more frequently than bottom-quartile individuals (Fig. 3c), confirming that MCMC-OCS correctly identifies which individuals to favour, even when precise contribution magnitudes vary across iterations. Using the **A** matrix for the same subset yielded modestly worse agreement across all metrics (Supplementary Table S8), while expanding to the full 5,525-tree population further reduced consistency (mean overlap: 14.1 [**H**] and 10.4 [**A**] individuals), reflecting greater within-family ranking ambiguity at larger sample sizes.

**Figure 3.**
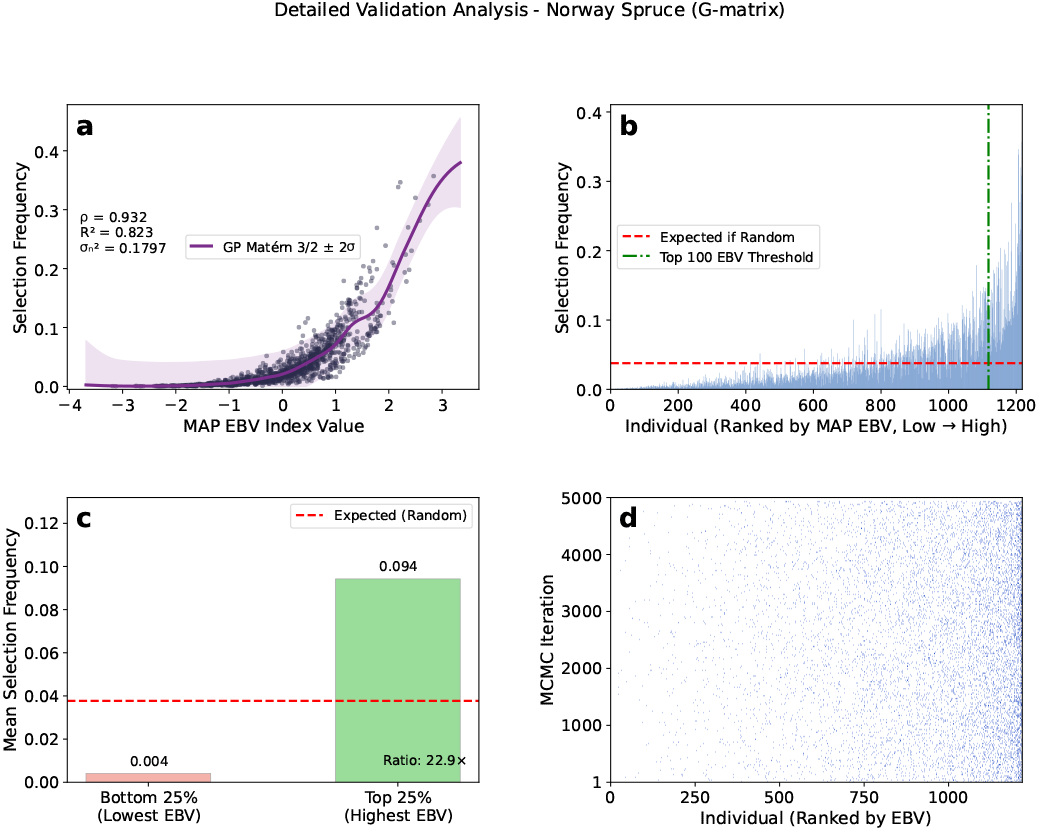
Detailed validation analysis of MCMC-based optimum contribution selection uncertainty for the Norway spruce breeding population (**G**-matrix, *n* = 1,218). **a** Relationship between MAP breeding values and selection frequency across MCMC iterations, with GP regression trend (Matérn 3/2 kernel) indicating a strong positive correlation despite high iteration-level variability. Annotation gives Spearman correlation (*ρ*), GP variance explained (*R*^2^), and inferred noise variance (*σ*^2^). **b** Selection frequency for individuals ranked by MAP breeding value, showing a clear gradient from low-EBV (rarely selected) to high-EBV (frequently selected) individuals; red dashed line = expected random selection frequency; green dash-dot = top-100 EBV threshold. **c** Quartile comparison of mean selection frequencies for the bottom vs. top 25% of breeding values; red dashed line = expected random frequency. **d** Binary heatmap of selection patterns across a sample of MCMC iterations (white = not selected, blue = selected with an average 46 of individuals per iteration), with individuals sorted by breeding value rank. Despite substantial iteration-level uncertainty (mean overlap 26.6 individuals), high-EBV individuals are consistently preferentially selected.

#### Robustness analysis of contributions

To motivate the CVaR-OCS approach, we first examined which MAP-OCS selections in Norway spruce are most vulnerable to EBV uncertainty using the individual robustness scores described in Materials and Methods. For the genotyped subpopulation (*n* = 1,218), 25 of 154 MAP-OCS selections were identified as high-risk (bottom-quartile robustness scores), including three of the top-25 contributors. Hence, high contribution under MAP EBVs does not guarantee stability under posterior uncertainty. Re-running OCS after excluding these individuals improved mean robustness by 16.5% at a genetic gain cost of only 3.0%, but paradoxically increased contribution concentration (Gini +5.6%, top10% concentration +12.9%), reflecting a structural limitation of post-hoc exclusion. Removing individuals forces the optimizer to concentrate contributions among the remaining high-EBV candidates, reintroducing fragility elsewhere in the selected population. This limitation directly motivates the suggested principled uncertainty-aware approach CVaR-OCS. Full robustness score results, risk classification figures, and contribution distribution metrics are provided in the Supplementary Material. Robustness score computation required approximately *n*_sel_ × *n*_MCMC_ = 88 × 200 = 17,600 MAP-OCS solves for the *n* = 1,218 subpopulation (*ca*. 1,133 min sequentially; *ca*. 57 min on 20 cores in parallel, see Supplementary Results), confirming that the diagnostic is practical when parallelised over candidates and/or after subsampling.

#### CVaR-OCS efficiency frontier and operating point selection

These results motivate a fundamentally different approach. Rather than identifying and removing uncertain individuals after optimisation, CVaR-OCS incorporates EBV uncertainty directly into the OCS objective, producing contribution vectors that are robust to the uncertainty quantified by the MCMC evaluation.

To evaluate the trade-off between expected genetic gain and resilience to EBV uncertainty, we applied CVaR-OCS to the Norway spruce genomic dataset (*n* = 1,218; Θ = 0.02) across three confidence levels (*α* ∈ { 0.90, 0.95, 0.99} ) and a range of CVaR weights (*µ* ∈ { 0.50, …, 10.00} ). For each (*α, µ*) combination, the CVaR-OCS objective penalises solutions whose expected genetic gain deteriorates under the worst (1 − *α*) × 100% of MCMC-sampled EBV realisations, thereby directing contributions away from individuals whose breeding values are sensitive to posterior uncertainty.

The resulting efficiency frontier (Figure 4a) exhibits a consistent concave shape at all *α* levels. Small increases in *µ* initially produce substantial improvements in CVaR_95_ at negligible cost to expected gain, with marginal returns declining rapidly beyond a species-specific threshold. To identify the optimal operating point for each *α*, we applied the geometric elbow criterion that locates the frontier point with maximum perpendicular distance from the chord connecting the MAP-OCS baseline to the maximum-*µ* solution (see *CVaR-OCS parameter selection* in Materials and Methods). Elbow points and their associated gain– robustness trade-offs are summarised in Table 3 and illustrated in Figure 4. The sharpest and most cost-efficient elbow was observed at *α* = 0.90, *µ* = 1.25, where marginal efficiency, measured as the ratio of CVaR_95_ improvement to genetic gain loss, dropped by 58% between *µ* = 1.25 and *µ* = 1.50 (from 1.84 to 0.77 ΔCVaR_95_%/Δgain loss%), confirming a pronounced change in the efficiency regime (Figure 4, panel b). More conservative confidence levels identified their elbows at *µ* = 1.75 (*α* = 0.95 and *α* = 0.99), but at gain costs of 2.06% and 3.67%, respectively: three to five times higher than the *α* = 0.90 elbow. We therefore adopted *α* = 0.90, *µ* = 1.25 as the recommended operating point for all subsequent analyses.

**Table 3.** Summary of CVaR-OCS efficiency frontier elbow points and the MAP-OCS baseline for Norway spruce (*Picea abies*; *n* = 1,218; Θ = 0.02). Elbow points were identified using a geometric perpendicular-distance criterion applied to the normalised gain– CVaR_95_ efficiency frontier (see Materials and Methods). Gain loss and CVaR_95_ improvement are expressed as percentages relative to the MAP-OCS baseline. The recommended operating point (*α* = 0.90, *µ* = 1.25) is shown in bold. Ad-hoc exclusion results are reported for directional comparison only (see text).

| Method | $\alpha$ | $\mu$ | $\bar{G}$ | Gain loss (%) | CVaR<sub>95</sub> | CVaR<sub>95</sub> imp. (%) | $n_{\text{sel}}$ | Gini |
| --- | --- | --- | --- | --- | --- | --- | --- | --- |
| MAP-OCS | — | — | 0.6436 | — | 0.4099 | — | 145 | 0.483 |
| **CVaR-OCS (elbow)** | **0.90** | **1.25** | **0.6391** | **−0.70** | **0.4369** | **+6.60** | **159** | **0.460** |
| CVaR-OCS (elbow) | 0.95 | 1.75 | 0.6304 | −2.06 | 0.4480 | +9.30 | 159 | 0.444 |
| CVaR-OCS (elbow) | 0.99 | 1.75 | 0.6200 | −3.67 | 0.4423 | +7.93 | 170 | 0.463 |
| Ad-hoc exclusion<sup>†</sup> | — | — | — | −4.35 | — | −3.70 | 118 | 0.510 |

<sup>†</sup>Ad-hoc exclusion: bottom-quartile MAP-OCS selections by robustness score ( $n = 40$ ) excluded; OCS re-run on remaining candidates. Absolute gain and CVaR<sub>95</sub> values are not directly comparable to CVaR-OCS due to differences in EBV standardisation between analysis pipelines; percentage changes relative to the respective MAP-OCS baseline are reported.

**Figure 4.**
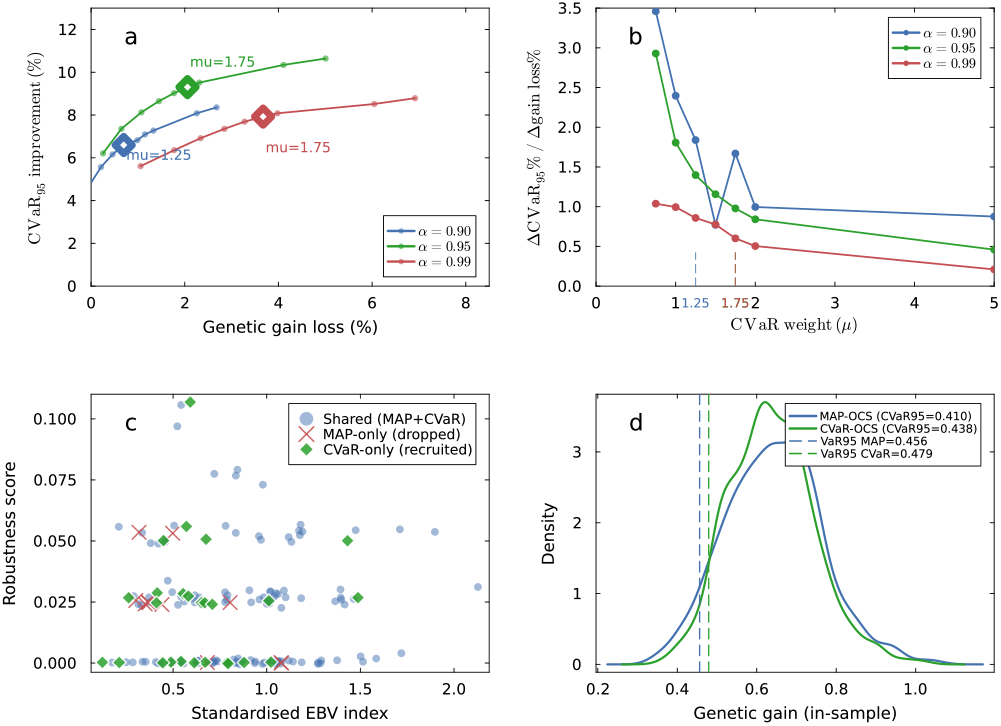
CVaR-OCS analysis for Norway spruce (*Picea abies*; *n* = 1,218 Θ = 0.02). **a** Gain–tail-risk efficiency frontier across three confidence levels (*α* ∈ { 0.90, 0.95, 0.99} ) and CVaR weights (*µ* ∈ {0.50, …, 10.00} ). Each curve shows the trade-off between genetic gain loss (%) and CVaR_95_ improvement (%) relative to the MAP-OCS baseline. Open diamonds mark the elbow point for each *α* level, identified by a geometric perpendicular-distance criterion applied to the normalised frontier (*CVaR-OCS parameter selection* in Materials and Methods). **b** Marginal efficiency of CVaR weighting — the ratio of CVaR_95_ improvement to genetic gain cost between successive *µ* values. A sharp drop in marginal efficiency confirms the recommended operating point at *α* = 0.90, *µ* = 1.25 (dashed vertical line); the x-axis is truncated at *µ* = 5 for clarity. **c** Standardised EBV index vs MCMC robustness score for all candidates at the recommended operating point, coloured by selection group: individuals retained by both MAP-OCS and CVaR-OCS (Shared; blue circles), MAP-OCS selections dropped by CVaR-OCS (MAP-only; red crosses), and new individuals recruited by CVaR-OCS (CVaR-only; green diamonds). Robustness score reflects the expected genetic gain loss when an individual is excluded across MCMC iterations. **d** Kernel density of genetic gain realised by MAP-OCS (blue) and CVaR-OCS (green) solutions evaluated across all 5,000 MCMC EBV scenarios. Dashed vertical lines mark the 5th-percentile gain (VaR_95_) for each solution; CVaR_95_ values (mean gain below the 5th percentile) are given in the legend. CVaR-OCS improves CVaR_95_ by 6.60% and VaR_95_ by 5.00% relative to MAP-OCS at a genetic gain cost of 0.70%, confirmed by a significantly reduced gain variance (F-test: *p* < 0.001) with no significant reduction in mean gain (t-test: *p* = 0.21).

#### Solution properties at the recommended operating point

At *α* = 0.90, *µ* = 1.25, CVaR-OCS achieved a CVaR_95_ improvement of 6.60% relative to MAP-OCS (0.410 → 0.437), alongside a 5.00% improvement in VaR_95_ (0.456 → 0.479), while reducing expected genetic gain by only 0.70% (0.644 → 0.639; Figure 4, panel c d). This represents a highly favourable trade-off given the substantial EBV uncertainty characteristic of Norway spruce genomic evaluations, where G-matrix-based correlations are considerably weaker and more variable than pedigree-based estimates due to the relatively modest genotyped sample size.

The CVaR-OCS solution selected 159 parents compared to 145 under MAP-OCS, a 9.7% broader selection base. Of these, 135 individuals were shared between the two solutions, while 10 MAP-OCS selections were dropped and 24 new individuals recruited. The Gini coefficient of contribution weights decreased from 0.483 (MAP-OCS) to 0.460 (CVaR-OCS), a 4.9% reduction indicating a modest redistribution toward more equitable parental contributions. Evaluated across all 5,000 MCMC scenarios, the gain distribution under CVaR-OCS was significantly less variable than under MAP-OCS (F-test: *p* < 0.001), with no significant difference in mean expected gain (t-test: *p* = 0.21), confirming that CVaR-OCS improves tail-gain security without penalising average performance. A single CVaR-OCS solve at the recommended operating point required 84.0 ± 4.4 s (mean ± SD, *N* = 10 replicates), compared with 3.9 ± 0.9 s for MAP-OCS, an approximately 22 × overhead attributable to the *l* = 5,000 auxiliary variables added by the CVaR formulation (Supplementary Results).

#### Comparison with ad-hoc robustness-based exclusion

As a reference, we evaluated an intuitive but theoretically ungrounded alternative. By identifying MAP-OCS selections whose robustness scores fell in the bottom quartile (*n* = 40 of 158 selected; mean robustness score = −0.005 vs. +0.039 for the retained group), excluding them from the candidate set, and re-running OCS on the remaining individuals. This approach incurred a genetic gain loss of 4.35%, more than six times the cost of CVaR-OCS at the recommended operating point, while paradoxically worsening CVaR_95_ by 3.7% relative to MAP-OCS. The deterioration reflects a structural limitation of post-hoc exclusion. Removing individuals forces the constrained OCS to concentrate contributions among the remaining high-EBV candidates, increasing sensitivity to posterior uncertainty in exactly those selections. CVaR-OCS avoids this by incorporating uncertainty directly into the optimisation objective, allowing it to simultaneously broaden the selection base. This reduces contribution concentration, and improve tail-gain security, at a fraction of the genetic gain cost.

## Discussion

Optimum contribution selection (OCS) is a widely used framework for balancing genetic gain and inbreeding in breeding and conservation programs (Meuwissen 1997; Hinrichs *et al*. 2006; Wellmann 2019; Mullin and Belotti 2015; Wang *et al*. 2017; Hjortø *et al*. 2022; El-Kassaby *et al*. 2024; Waldmann 2025), yet the impact of uncertainty in estimated breeding values (EBVs) on optimized contributions has received comparatively little attention. Here we address this gap through two complementary contributions. First, we introduce CVaR-OCS, a single-solve uncertainty-aware formulation that directly incorporates the joint posterior distribution of EBVs into the optimization objective via the Conditional Value at Risk (CVaR) risk measure. Second, we use Monte Carlo sampling across MCMC iterations (MCMC-OCS) as a diagnostic tool that exposes the instability of MAP-OCS decisions under EBV uncertainty, and we develop individual-level robustness scores to identify high-risk selections. CVaR-OCS is the principal methodological advance: rather than solving one OCS problem per posterior draw and aggregating the results post hoc, it solves a single quadratic program that explicitly optimizes expected genetic gain subject to a formal tail-risk constraint, providing a principled and computationally efficient path to uncertainty-aware breeding decisions.

The choice of CVaR as the risk measure for OCS rests on solid theoretical foundations from financial portfolio optimization. CVaR, also known as Expected Shortfall, is a coherent risk measure in the sense of Artzner *et al*. (1999). It satisfies subadditivity, monotonicity, translation invariance, and positive homogeneity. Subadditivity in particular is essential for a diversification-based framework such as OCS, because it guarantees that combining two contribution portfolios cannot increase tail risk beyond the sum of their individual risks. Crucially, CVaR admits a tractable quadratic programming reformulation (Rockafellar and Uryasev 2000, 2002), which is what makes CVaR-OCS practical: the auxiliary-variable representation transforms the otherwise non-smooth tail expectation into a QP with *l* additional linear constraints, one per MCMC scenario, that can be solved directly with a standard interior-point or ADMM solver such as COSMO (Garstka *et al*. 2019).

Given the relatively tight relatedness constraint (Θ = 0.02 in the Norway spruce case study), much of the genetic gain captured by OCS derives from exploiting within-family variation. However, when EBV uncertainty is propagated through the posterior distribution, within-family rankings shift dramatically across MCMC iterations compared to the MAP-optimal solution. These instabilities are directly connected to the role of Mendelian segregation variance in OCS (Avendaño *et al*. 2004; Villanueva *et al*. 2006; Woolliams *et al*. 2015; Howard *et al*. 2018). Avendaño *et al*. (2004) showed that optimized contributions are allocated according to each candidate’s Mendelian sampling term, that is, their superiority over the parental average. Thus, accurate estimation of this within-family additive component is critical for OCS to deliver its theoretical advantages. Howard *et al*. (2018) confirmed this in a commercial pig population. Our framework makes the consequences of imprecise Mendelian sampling estimates explicit. Individuals whose within-family advantage is poorly estimated receive highly variable contributions across posterior draws, and are precisely the individuals flagged as high-risk by the MCMC robustness scores (Supplementary Material). Strategies to improve within-family precision include broader progeny testing (Kerr *et al*. 2015), increasing family size and representation (Chu *et al*. 2019), and enhanced phenotyping protocols (Angidi *et al*. 2025).

Alongside CVaR-OCS, we developed two complementary descriptive metrics, individual robustness scores and the Gini coefficient of the contribution distribution, for characterising EBV uncertainty at the individual and population level respectively. The population-level metrics connect directly to the theory of long-term genetic contributions formalised by Woolliams *et al*. (1999) and reviewed by Woolliams *et al*. (2015): under the genetic contributions framework, the rate of inbreeding is proportional to the sum of squared long-term genetic contributions 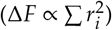, so the distribution of parental contributions determines not only single-generation genetic gain but also the long-run effective population size and inbreeding trajectory (Woolliams *et al*. 1999; Bijma 2012). A skewed contribution distribution (high Gini, high top-10% concentration) implies a narrow effective gene pool that accelerates genetic drift and inbreeding accumulation across generations, even when the single-generation coancestry constraint Θ is satisfied (Meuwissen *et al*. 2020). Monitoring the Gini coefficient and concentration ratio alongside CVaR and genetic gain therefore provides a window onto the multi-generational consequences of a given contribution vector, motivating their inclusion as diagnostic complements to the single-generation CVaR objective.

It is important to note that a low individual robustness score can reflect either high EBV uncertainty for that individual across MCMC iterations, or the availability of near-equivalent genetic substitutes in the candidate pool, or both. The score therefore quantifies replaceability under uncertainty rather than EBV uncertainty in isolation. This distinction is practically relevant in populations with structured family groups, where within-family Mendelian sampling variation means that siblings may share nearly identical EBVs under limited data (Woolliams *et al*. 1999; Avendaño *et al*. 2004). An individual with a highly uncertain EBV but no close genetic substitutes may still receive a high robustness score, whereas an individual with a stable EBV but several near-clones in the candidate set may score low. CVaR-OCS does not share this ambiguity, as it optimises directly against the joint posterior uncertainty of the full candidate pool simultaneously.

In Norway spruce, the dual metrics revealed a notable dissociation. Individual robustness improved after post-hoc exclusion of high-risk selections, but contribution concentration increased (Gini +5.6%, top-10% concentration +12.9%), indicating that population-level diversification did not co-move with individual-level stability. The mechanism is structural by construction, as OCS concentrates contributions on the highest-EBV individuals subject to the coancestry constraint (Meuwissen 1997). Removing candidates forces the optimiser to concentrate even more heavily among the remaining elite, reintroducing a different form of fragility elsewhere in the portfolio and, from the perspective of long-term contributions (Woolliams *et al*. 1999, 2015), narrowing the effective gene pool relative to the MAP-OCS baseline. This contrasts with the QTL-MAS results, where CVaR-OCS simultaneously broadened the selection base, reduced contribution concentration, and improved tail-gain security, i.e. a concordance that likely reflects the less tight coancestry constraint and larger candidate pool in that dataset, which give the optimiser more room to redistribute contributions without sacrificing gain (Gautason *et al*. 2023). The divergent behaviour between datasets underscores that individual robustness and portfolio concentration are not interchangeable metrics and that post-hoc exclusion cannot in general guarantee improvement on both dimensions simultaneously (Woolliams *et al*. 2015; Meuwissen *et al*. 2020). Notably, the robustness scores also revealed that several high-contributing individuals under MAP-OCS were simultaneously classified as high-risk (Supplementary Figure S7b), illustrating that a large MAP contribution does not imply stability under posterior uncertainty: an individual can dominate the MAP solution because their point-estimate EBV is high relative to the coancestry constraint, while their full posterior distribution overlaps substantially with competing candidates, making them replaceable across many MCMC iterations. This is the class of selections that both robustness scores and CVaR-OCS are designed to identify and penalise respectively.

The bottom-quartile threshold used to classify high-risk selections is illustrative rather than prescriptive. In practice, the threshold could be guided by: (i) a tolerable budget of genetic gain loss, framed within the classical gain–inbreeding trade-off of OCS (Meuwissen 1997); (ii) a natural break in the robustness score distribution, which often separates a small cluster of markedly low-scoring individuals from the remaining candidates; or (iii) the CVaR-OCS efficiency frontier itself, which provides a principled and continuous parameterisation of the robustness–gain trade-off through *α* and *µ* without requiring an arbitrary exclusion threshold. We recommend CVaR-OCS as the primary decision-making tool, with robustness scores serving as complementary diagnostics to identify which MAP-OCS selections are most exposed to posterior uncertainty. Further simulation work across multiple generations is needed to establish how the Gini coefficient and robustness scores behave over time and which monitoring statistics are most informative for long-term operational decision-making (Woolliams *et al*. 2015; Meuwissen *et al*. 2020).

The computational demands of the three framework components differ substantially and should guide implementation choices in practice. A single MAP-OCS solve is fast (3.9 ± 0.9 s on the Norway spruce genomic dataset), and CVaR-OCS adds a manageable overhead (84.0 ± 4.4 s; approximately 22× MAPOCS) arising from the *l* auxiliary variables introduced by the CVaR reformulation. This is a fixed cost that does not grow with repeated optimisation across scenarios, since CVaR-OCS solves only once regardless of chain length. We consider this overhead justified by the gain in robustness, the elimination of the arbitrary exclusion threshold, and the principled probabilistic interpretation of the CVaR objective; for breeding programs already investing in MCMC genetic evaluation, the additional computation is modest relative to the evaluation step itself. Robustness score calculation is substantially more demanding, scaling as *n*_sel_ × *n*_MCMC_ sequential OCS solves. In the Norway spruce scenario this amounted to approximately 1,133 minutes sequentially, reduced to around 57 minutes on 20 cores after subsampling to 200 MCMC draws (see Results). For large populations or dense genomic relationship matrices, exhaustive robustness score computation will often be impractical, and we recommend restricting it to situations where populations are small, pedigree-based matrices are available (reducing single-solve time by 4–18 fold relative to **G**; Supplementary Table S9), or where targeted diagnostic information about specific high-value candidates is operationally warranted. For routine uncertainty-aware OCS, CVaR-OCS is the recommended approach precisely because it delivers the robustness benefits of the full MCMC posterior in a single computationally efficient solve.

A recurring practical question in genomic OCS is whether the relationship matrix used in the coancestry constraint must be the same as the one used in the genetic evaluation step to infer breeding values, and if not, which matrix is preferable in each role. In principle, these two uses serve distinct purposes. The evaluation matrix determines the statistical model for decomposing phenotypic variance and thus affects EBV accuracy, while the OCS matrix controls which notion of relatedness is penalised in the coancestry constraint and therefore shapes the long-term inbreeding trajectory. There is no theoretical requirement that these two matrices be identical (Meuwissen *et al*. 2020), and empirical evidence on their relative merits in the OCS step remains mixed. Henryon *et al*. (2019) found that pedigree-based OCS (POCS) realised 14–34% more true genetic gain than genomic OCS (GOCS) under ABLUP, and 1.5–5.7% more under GBLUP, when both were constrained to the same rate of true inbreeding on the IBD scale. They attributed this to GOCS unnecessarily penalising allele frequency changes at markers driven by genetic drift, thereby restricting beneficial selection at QTL. By contrast, Gautason *et al*. (2023) found that when the number of sires is fixed and each approach is constrained on its own kinship scale, genomic OCS using VanRaden’s method 1 with base-population allele frequencies realised more genetic gain for a given level of kinship and inbreeding than pedigree-based OCS, because genomic relationships account for Mendelian sampling variation within families that pedigree relationships cannot resolve. The key difference between these two results is the comparison basis. Constraining POCS and GOCS to the same inbreeding rate on the same scale favours pedigree relationships, whereas allowing each approach to operate on its own scale and fixing operational constraints such as number of sires favours genomic relationships. Applying the same threshold Θ to **G** and **A** implies different effective inbreeding targets because **G**_*ij*_ includes a baseline IBS component reflecting founder allele frequencies, whereas **A**_*ij*_ measures IBD relative to a defined pedigree base (Hill and Weir 2011). In our Norway spruce analyses, this effect was modest (baseline average coancestry 0.012 genomic vs. 0.014 pedigree), but would become more consequential at tighter constraints or in more inbred populations, and practitioners should calibrate Θ to the matrix being used rather than applying a universal threshold across matrix types.

Our work builds on two key recent contributions to uncertainty-aware OCS. Pocrnic *et al*. (2023) demonstrated the value of propagating posterior EBV distributions through OCS using a simulated population, providing the conceptual foundation for the MCMC-OCS diagnostic procedure used here. Fogg *et al*. (2024) developed robust OCS using standard errors of EBVs, offering a practical solution applicable to frequentist inference settings. CVaR-OCS is complementary to both: it extends the posterior sampling idea of Pocrnic *et al*. (2023) into a single unified optimization problem, and it subsumes the minimax robust OCS of Fogg *et al*. (2024) as a limiting case. As the CVaR confidence level *α* → 1, CVaR-OCS converges to a worst-case formulation equivalent to SE-based robust OCS (Fogg *et al*. 2024). A further distinction is that CVaR-OCS propagates the full joint posterior covariance structure of breeding values implicitly through the scenario set, whereas SE-based approaches treat each candidate’s uncertainty independently. This joint propagation is particularly relevant in populations with strong family structure or high genomic relatedness among elite candidates. The QTL-MAS simulation results provide an initial empirical validation. CVaR-OCS recovered one additional oracle-optimal individual compared to MAP-OCS and reduced the variance of the gain distribution, consistent with the theoretical expectation that tail-risk protection should move the solution closer to a robustly-optimal portfolio. However, the improvement in Jaccard similarity to the oracle was modest (+11%), and these results derive from a single-generation benchmark with a relatively small candidate population. Multi-generation simulation studies comparing CVaR-OCS and MAP-OCS under known true breeding values are needed to characterize the long-term effects of incorporating EBV uncertainty on genetic gain trajectories and inbreeding accumulation, and to determine whether the performance advantage of CVaR-OCS grows, diminishes, or remains stable across generations. Together with the independent contributions of Fogg *et al*. (2024) and Pocrnic *et al*. (2023), these converging efforts underscore the growing recognition that EBV uncertainty is an important and tractable dimension of OCS that the field is only beginning to address systematically.

## Data availability

Norway spruce phenotypic and genotypic data with recoded IDs are provided here: https://github.com/jonhar97/MCMC_uncertainty_OCS/.

### Acknowledgments

This work was supported by grant from Föreningen Skogsträdsförädling project ID 23-511. The authors thank the Associate Editor Dr Jeffrey Endelmann and three anonymous reviewers for their constructive comments, which substantially improved the manuscript. The online version contains supplementary material here;

