## Supplementary material for "Uncertainty-aware breeding decisions: MCMC-based optimum contribution selection increases breeding decision robustness"

#### Abstract

This supplementary material provides full methodological details and results for the MCMC-based robustness score analysis and contribution distribution metrics described in the main manuscript, together with optimization solver benchmarks. Individual robustness scores identified 25 high-risk MAP-OCS selections in Norway spruce; post-hoc exclusion improved individual robustness but increased contribution concentration in Norway spruce, motivating the CVaR-OCS approach developed in the main manuscript.

**Keywords:** Estimated breeding values; Posterior uncertainty; Genetic evaluation; Selection optimization; Norway spruce

#### Materials and Methods

##### Comparison of four different optimization packages

***Ipopt*** The advent of efficient interior point optimization methods has enabled the tractable solution of large-scale linear and nonlinear programming problems. The standard for interior point optimization is Ipopt, originally developed by Wächter and Biegler (2006). Ipopt transforms inequality constraints into logarithmic barrier terms in the objective and solves a sequence of equality constrained problems for decreasing values of the barrier parameter. For each barrier subproblem, a modified Newton algorithm with line search is used to obtain a solution to the problem. Each step of the Newton algorithm requires potentially multiple solutions to perturbed sparse symmetric indefinite systems of linear equations. Because of this inner algorithmic dependency, the performance of Ipopt is crucially dependent on the properties and performance of the selected linear solver that defaults to MUMPS in Julia. The computing time of Ipopt also relies on the external computation of the first and second derivatives, which is accomplished by the modeling language interface that in our case is provided by JuMP. We have used the Ipopt.jl library in Julia because it can solve large convex quadratic optimization problems (Wächter and Biegler 2006).

***ECOS*** The embedded conic solver (ECOS) is also an interior point solver for SOCP designed specifically for embedded applications. The main interior point algorithm is a standard primal-dual Mehrotra predictor-corrector method with Nesterov-Todd scaling and self-dual embedding, with search directions found via a symmetric indefinite Karush-Kuhn-Tucker system, chosen to allow stable factorization with a fixed pivoting order. ECOS is written in low-level, single-threaded, library-free ANSI-C which in Julia is available through the ECOS.jl package (Domahidi *et al.* 2013).

***SCS*** The Splitting Conic Solver (SCS) is based on a first-order quadratic cone programming (QCP) algorithm that is designed to efficiently handle large problem sizes and provide moderately accurate solutions. The algorithm uses Douglas-Rachford (DR) splitting to a homogeneous embedding of the linear complementarity problem (LCR). DR splitting is equivalent to ADMM under a particular change of variables and both are instantiations of the proximal point method (Parikh and Boyd 2014). The algorithm's per-iteration cost is almost identical to the linear-convex case and to applying the splitting method directly to the original problem. The homogeneous embedding approach offers advantages over methods that generate certificates of infeasibility based on diverging sequences. Specifically, infeasibility certificates are generated by convergence, providing more flexibility in achieving a solution, such as using inexact or stochastic updates. Numerical evidence suggests that the homogeneous embedding approach can converge to a solution slightly faster than direct approaches. The algorithm can solve convex quadratic cone programs involving any combination of nonnegative, second-order, semidefinite, exponential, and power cones. The algorithm has been implemented in C and is freely available in the solver SCS v3.0 via the Julia package SCS.jl (O'Donoghue 2021).

***COSMO*** The Conic Operator Splitting Method (COSMO) solver is based on an operator splitting algorithm that can handle convex optimization problems with quadratic objective function and conic constraints. At each step, the ADMM algorithm alternates between solving a quasi-definite linear system with a constant coefficient matrix and a projection onto convex sets. The low per-iteration computational cost makes the method particularly efficient for large problems, e.g. SDPs that arise in portfolio optimization which is closely related to our OCS problem. Moreover, the COSMO solver uses chordal decomposition techniques and a new clique merging algorithm to effectively exploit sparsity in large, structured SDPs. COSMO is implemented in pure Julia and available in the package COSMO.jl (Garstka *et al.* 2019).

#### MCMC chain length and convergence diagnostics

Genetic parameters and individual estimated breeding values (EBVs) were obtained using Bayesian multivariate GBLUP models implemented in JWAS.jl (Cheng *et al.* 2018). A total of 5,000 posterior samples were retained after discarding a burn-in of 2,000 iterations, with one sample saved per 10 iterations (total chain length 52,000). Note that JWAS does not implement a `thinning` argument; the sampling interval is controlled via `output_samples_frequency`, which governs both the thinning rate and the writing of posterior samples to disk.

Convergence was assessed by visual inspection of trace plots and by computing the effective sample size (ESS) for all heritability and genetic (co)variance parameters using the `MCMCDiagnosticTools.jl` package (Kumar *et al.* 2022). ESS quantifies the number of effectively independent draws in a correlated chain; values above 200 are generally considered adequate for posterior inference (Gelman *et al.* 2013).

For the primary genomic analysis (**G**-matrix, $n = 1{,}218$), all heritability ESS values exceeded 627 (ESS/n $\geq$ 0.125; Table S1), and trace plots showed stable mixing around stationary means throughout the chain (Figures S1–S2). For the blended relationship matrix analysis (**H**-matrix, $n = 5{,}525$), ESS values for the two continuous growth traits (Hjd17, Htv17) exceeded 200, while the low-heritability traits Sprant17 ($h^2 \approx 0.063$) and Lev17 ($h^2 \approx 0.057$, threshold model) showed ESS in the range 55–69 (Figures S3–S4). Low ESS for low-heritability parameters is a known feature of GBLUP chains, arising from a flat likelihood surface that increases autocorrelation; it does not indicate non-stationarity, as confirmed by the trace plots. A longer chain (chain_length $=$ 300,000; output_samples_frequency $=$ 50, yielding $\approx$ 5,960 retained samples) has been initiated for the H-matrix model and results will be updated accordingly. The GBLUP formulation involves substantially fewer free variance parameters than whole-genome marker-effect models, and the autocorrelation structure is accordingly more favourable (Sorensen and Gianola 2002).

**Table S1** Effective sample size (ESS) for heritability parameters from MCMC chains (5,000 retained samples after burn-in and thinning). ESS was computed using `MCMCDiagnosticTools.jl`. G-matrix: $n = 1{,}218$ genotyped individuals, three traits. H-matrix: $n = 5{,}525$ individuals, four traits (Lev17 fitted as threshold model).

| Matrix | Trait | ESS | ESS/n |
|---|---|---|---|
| G ($n = 1{,}218$) | Hjd17 | 765 | 0.153 |
| | Htv17 | 773 | 0.155 |
| | Sprant17 | 627 | 0.125 |
| H ($n = 5{,}525$) | Hjd17 | 216 | 0.043 |
| | Htv17 | 226 | 0.045 |
| | Sprant17 | 69 | 0.014 |
| | Lev17 | 66 | 0.013 |

*Low ESS reflects flat likelihood surface for low-heritability traits ($h^2 < 0.07$); trace plots confirm stationarity. Extended chain in progress.

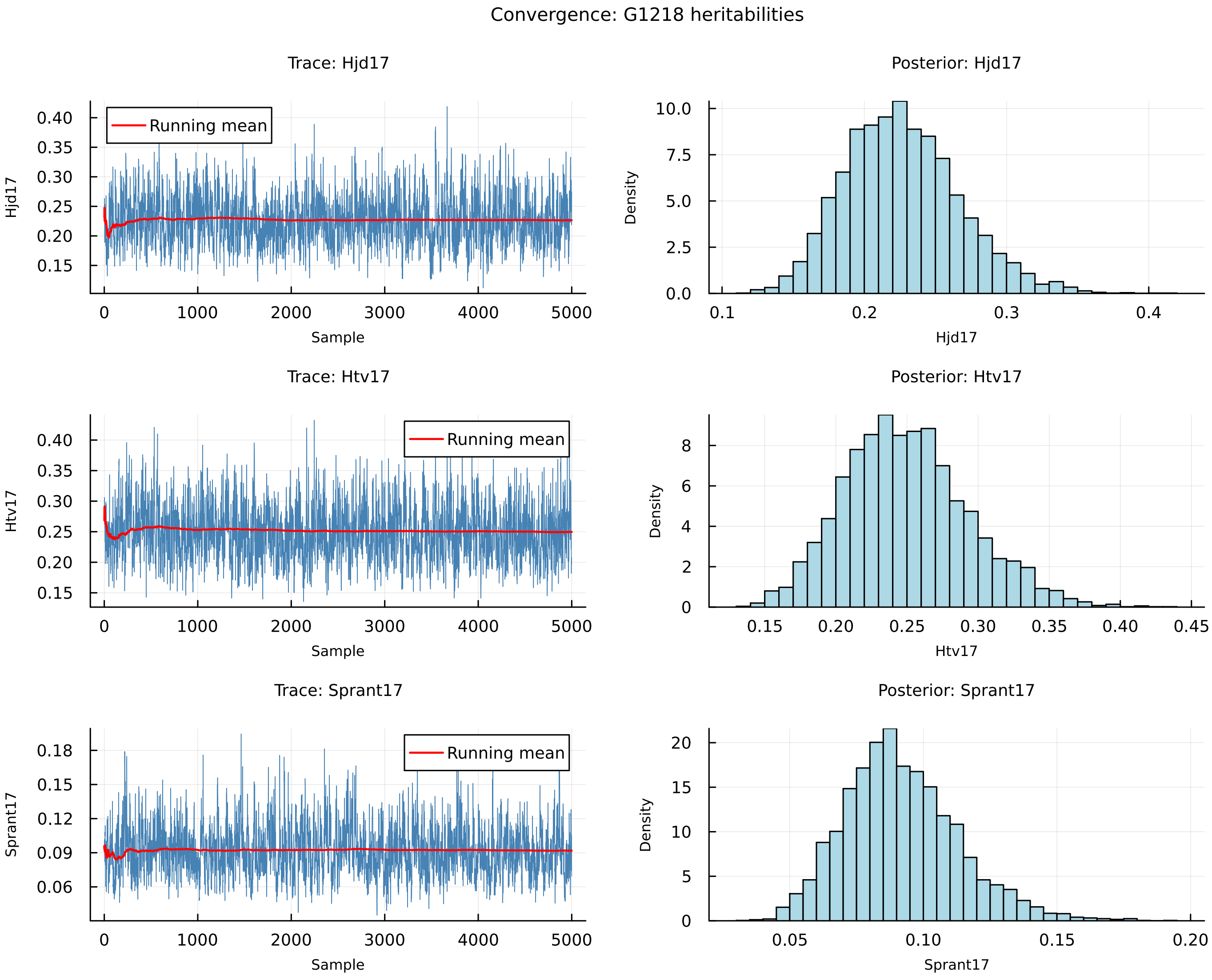


**Figure S1** MCMC convergence diagnostics for heritability parameters — G-matrix model ($n$ = 1,218, three traits). Left panels: trace plots with running mean (red line) showing stable mixing throughout the chain (5,000 retained samples after burn-in of 2,000 and thinning by 10; total chain length 52,000). Right panels: posterior marginal distributions. Effective sample sizes: Hjd17 = 765, Htv17 = 773, Sprant17 = 627 (all ESS/n $\geq$ 0.125).

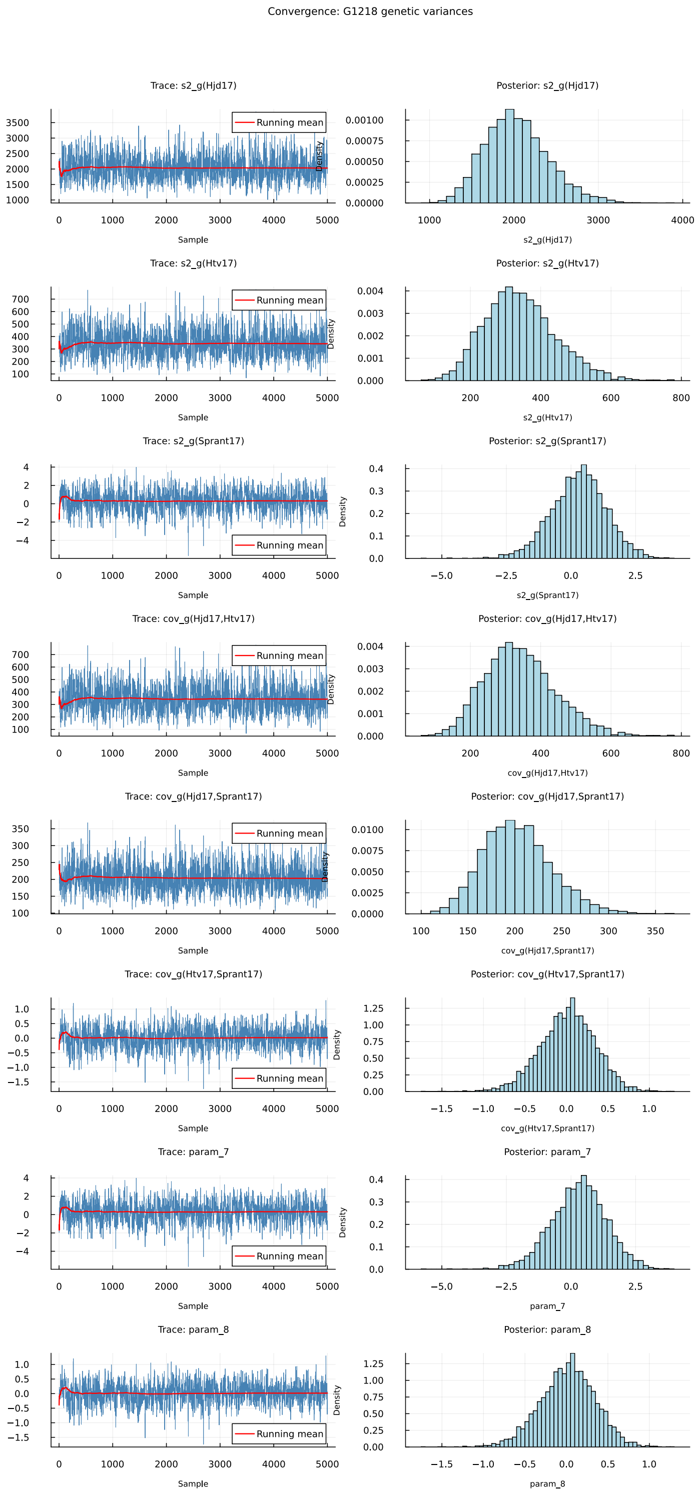


**Figure S2** MCMC convergence diagnostics for genetic variance and covariance parameters — G-matrix model ($n = 1{,}218$, three traits). Left panels: trace plots with running mean (red line). Right panels: posterior marginal distributions. All ESS $\geq$ 618 (ESS/n $\geq$ 0.124). Parameters labelled `param_7`–`param_9` correspond to lower-triangle elements of the symmetric genetic covariance matrix written redundantly by JWAS and are not separately interpreted.

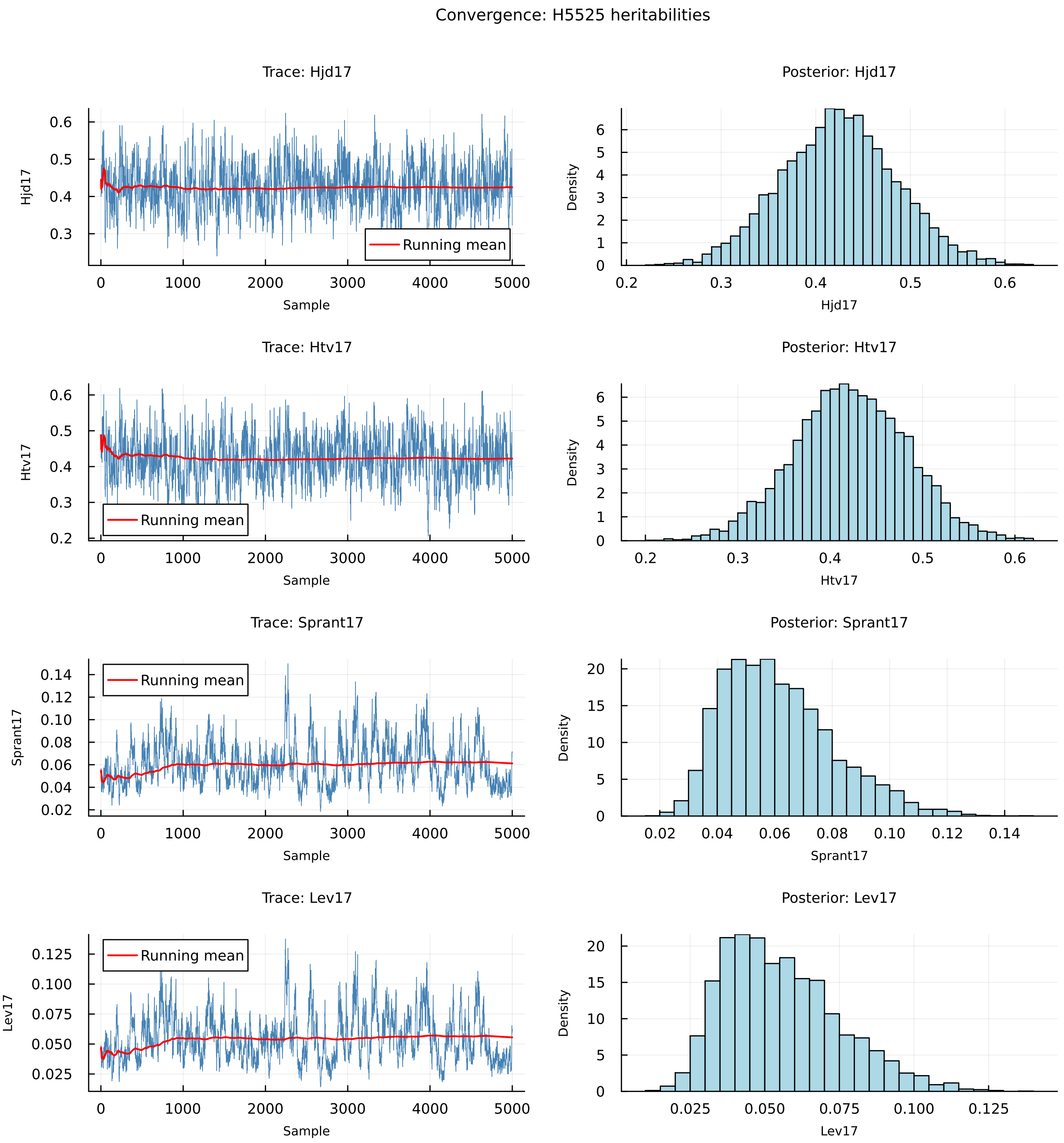


**Figure S3** MCMC convergence diagnostics for heritability parameters — H-matrix model ($n = 5{,}525$, four traits). Left panels: trace plots with running mean (red line). Right panels: posterior marginal distributions. Effective sample sizes: Hjd17 = 216, Htv17 = 226, Sprant17 = 69, Lev17 = 66 (5,000 retained samples; see Table S1). Low ESS for Sprant17 and Lev17 reflects the flat likelihood surface for low-heritability traits ($h^2 < 0.07$); trace plots confirm stationarity. *Note: figure will be updated following completion of extended chain (chain_length = 300,000).*

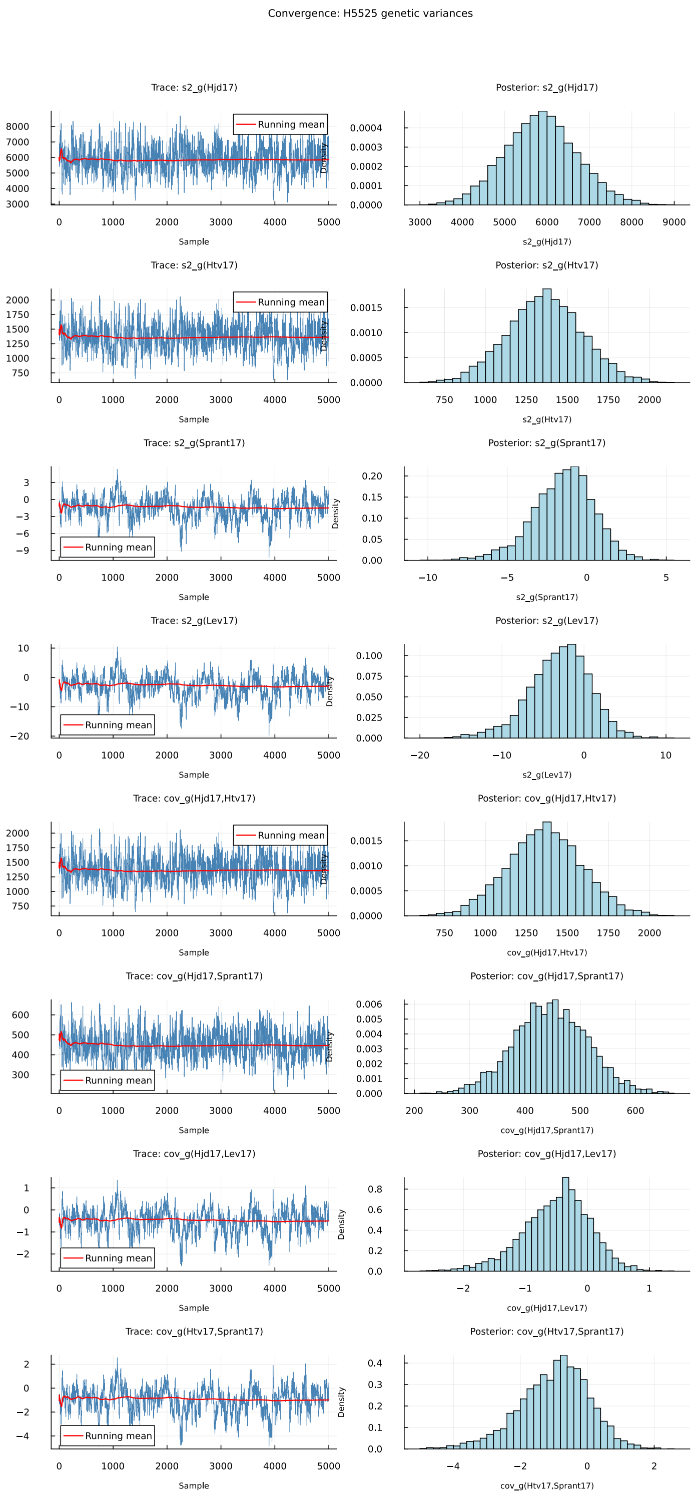


**Figure S4** MCMC convergence diagnostics for genetic variance and covariance parameters — H-matrix model ($n = 5{,}525$, four traits). Left panels: trace plots with running mean (red line). Right panels: posterior marginal distributions. ESS values for Sprant17 and Lev17 variance components are in the range 54–69 (ESS/n $\approx$ 0.011–0.014), consistent with the low heritability of these traits; all trace plots show stable behaviour around stationary means. *Note: figure will be updated following completion of extended chain (chain_length* $=$ 300,000*).*

#### MCMC-based robustness scores and contribution distribution metrics

##### Individual robustness scores

To quantify the stability of MAP-OCS selections across the posterior distribution, we defined a robustness score measuring each individual's importance to the breeding program as the expected genetic gain loss when that individual is excluded from the candidate set. For each individual $i$ and MCMC iteration $j$:

1. **Baseline genetic gain.** Calculate expected genetic gain using the complete breeding population:

$$G_{\text{baseline},j} = \sum_{k=1}^{n} \text{BV}_{k,j} \times c_{k,j}, \tag{1}$$

   where $\text{BV}_{k,j}$ is the standardised breeding value of individual $k$ in iteration $j$ and $c_{k,j}$ is the corresponding contribution. Breeding values were standardised within each MCMC iteration before running OCS to avoid weighting differences across the MCMC procedure.

2. **Modified genetic gain.** Set individual $i$'s breeding value to a value below the minimum (effectively removing them), re-run OCS, and calculate the new expected genetic gain:

$$G_{\text{modified},j} = \sum_{k=1}^{n} \text{BV}_{k,j} \times c'_k, \tag{2}$$

   where $c'_k$ represents the new optimal contributions excluding individual $i$.

3. **Individual impact:**

$$\Delta G_{i,j} = G_{\text{baseline},j} - G_{\text{modified},j}. \tag{3}$$

4. **Mean robustness score:**

$$S_i = \frac{1}{l} \sum_{j=1}^{l} \Delta G_{i,j}. \tag{4}$$

A stable robustness metric expressing robustness as a percentage of baseline genetic gain was also calculated:

$$P_i = \frac{S_i}{\bar{G}} \times 100, \tag{5}$$

where $\bar{G} = l^{-1} \sum_{j=1}^{l} G_{\text{baseline},j}$. Higher robustness scores indicate individuals whose removal causes substantial loss of genetic gain, while lower scores indicate individuals whose contribution is less critical or easily replaceable. The robustness score thus provides an uncertainty-aware measure of individual importance that accounts for variability in EBV estimates across the posterior distribution. To further speed up computation, 100 out of 1,000 MCMC samples were randomly selected for robustness score calculations.

##### Marginal gain distribution metrics

To complement individual robustness scores with population-level risk metrics — conceptually similar to portfolio diversification measures in finance — we quantified the distribution of genetic contributions across selected individuals. For each OCS solution, the marginal gain of individual $i$ was calculated as

$$M_i = c_i \times \text{BV}_i, \tag{6}$$

where $c_i$ is the optimal contribution and $\text{BV}_i$ is the standardised breeding value of individual $i$, representing the expected genetic gain attributable to that individual. Contribution distribution structure was assessed using four complementary metrics:

1. **Mean marginal gain:**

$$\bar{M} = \frac{1}{n_{\text{sel}}} \sum_{i \in \mathcal{S}} M_i, \tag{7}$$

   where $\mathcal{S}$ is the set of selected individuals ($c_i > 10^{-4}$) and $n_{\text{sel}} = |\mathcal{S}|$. Lower mean marginal gains indicate more evenly distributed breeding populations.

2. **Maximum marginal gain:**

$$M_{\max} = \max_{i \in \mathcal{S}} M_i. \tag{8}$$

   This quantifies the vulnerability to the loss of the most critical individual.

3. **Concentration ratio:**

$$\text{Conc}_{10} = \frac{\sum_{i \in \mathcal{T}_{10}} M_i}{\sum_{i \in \mathcal{S}} M_i} \times 100, \tag{9}$$

   where $\mathcal{T}_{10}$ represents the top 10% of individuals ranked by contribution.

**Table S2** Phenotypic data summary for Norway spruce (*Picea abies*) across both analysis datasets. The full pedigree dataset ($n = 5{,}525$) was used with the blended **H**-matrix and pedigree **A**-matrix models; the genotyped subset ($n = 1{,}218$) was used with the genomic **G**-matrix model. Both datasets span two field trials (S23F8820476: 2,401/568 trees; S23F8820477: 3,124/650 trees) established at the same sites. The genotyped subset is a strict subset of the full pedigree dataset (confirmed by individual ID matching).

| Dataset | Trait | Records ($n$) | Missing (%) | Mean | SD | Type |
|---|---|---|---|---|---|---|
| Full pedigree $n = 5{,}525$ 142 families | Hjd17 | 4,462 | 19.2 | 411.6 | 174.7 | Continuous |
| | Htv17 | 4,319 | 21.8 | 107.5 | 57.2 | Continuous |
| | Sprant17 | 4,462 | 19.2 | 0.2 | 0.5 | Continuous |
| | Lev17 | 5,525 | 0.0 | 0.8 | 0.4 | Threshold[a] |
| Genotyped subset $n = 1{,}218$ 126 families | Hjd17 | 1,217 | 0.1 | 458.5 | 153.8 | Continuous |
| | Htv17 | 1,214 | 0.3 | 117.5 | 55.3 | Continuous |
| | Sprant17 | 1,217 | 0.1 | 0.2 | 0.5 | Continuous |

Hjd17: dominant height at age 17 (cm); Htv17: stem volume at age 17 ($dm^3$); Sprant17: ramicorn count at age 17; Lev17: survival at age 17 (binary: 0 = dead, 1 = alive). Missing values for continuous traits arise from mortality prior to measurement. Lev17 has no missing values as survival status was recorded for all individuals.

[a] Lev17 was excluded from the G-matrix model because only 1 of 1,218 genotyped trees died before age 17 (survival rate 99.9%), providing insufficient variation to estimate a genetic variance component. Survival rate in the full pedigree dataset was 80.8% (4,462 survivors of 5,525 trees).

**Table S3** Heritability estimates for Norway spruce (*Picea abies*) traits across relationship matrices

| Trait | G Matrix | H Matrix | A Matrix |
|---|---|---|---|
| Height growth, years 15–17 (Htv17) | 0.251 ± 0.048 | 0.442 ± 0.045 | 0.376 ± 0.069 |
| Height at 17 years, July (Hjd17) | 0.221 ± 0.056 | 0.442 ± 0.040 | 0.342 ± 0.049 |
| Height at 7 years (Hjd7) | 0.199 ± 0.081 | 0.341 ± 0.053 | 0.262 ± 0.058 |
| Ramicorns/spike knots (Sprant17) | 0.094 ± 0.040 | 0.108 ± 0.070 | 0.159 ± 0.091 |
| Survival/vitality (Lev17) | N/A[a] | 0.074 ± 0.023 | 0.121 ± 0.044 |

**G** matrix: Genomic relationships ($n = 1{,}218$); **H** matrix: Combined genomic and pedigree ($n = 5{,}525$); **A** matrix: Pedigree only ($n = 5{,}525$).

[a] No variation in survival among genotyped trees.

4. **Gini coefficient:** For $n_{\text{sel}}$ selected individuals with marginal gains sorted in ascending order ($M_{(1)} \leq M_{(2)} \leq \cdots \leq M_{(n_{\text{sel}})}$):

$$\text{Gini} = \frac{\sum_{i=1}^{n_{\text{sel}}} (2i - n_{\text{sel}} - 1)\, M_{(i)}}{n_{\text{sel}} \sum_{i=1}^{n_{\text{sel}}} M_{(i)}}, \tag{10}$$

where $\text{Gini} \in [0, 1]$; values near 0 indicate equal distribution and values near 1 indicate high concentration.

Together, these metrics provide complementary descriptive insight into how contribution decisions are distributed across the selected cohort under EBV uncertainty.

### Results

#### Genetic parameter estimates

Heritability and genetic correlation estimates across relationship matrices are presented in Tables S3–S4 and illustrated in Figures S5–S6. Phenotypic data summary statistics for both datasets are provided in Table S2.

**Table S4** Genetic correlation estimates between trait categories in Norway spruce (*Picea abies*) across relationship matrices

| Correlation Type | G Matrix | H Matrix | A Matrix |
|---|---|---|---|
| Growth–Growth[a] | 0.43 ± 0.22 | 0.78 ± 0.10 | 0.72 ± 0.13 |
| Growth–Ramicorns[b] | 0.08 ± 0.07 | −0.27 ± 0.09 | −0.06 ± 0.05 |
| Growth–Survival[c] | – | −0.28 ± 0.08 | −0.10 ± 0.06 |
| Ramicorns–Survival | – | 0.85 ± 0.34 | 0.75 ± 0.42 |

[a] Height traits: Hjd7, Hjd17, Htv17 correlations.
[b] Height traits vs Sprant17 correlations.
[c] Height traits vs Lev17 correlations. –: Not applicable due to trait absence in the respective model.

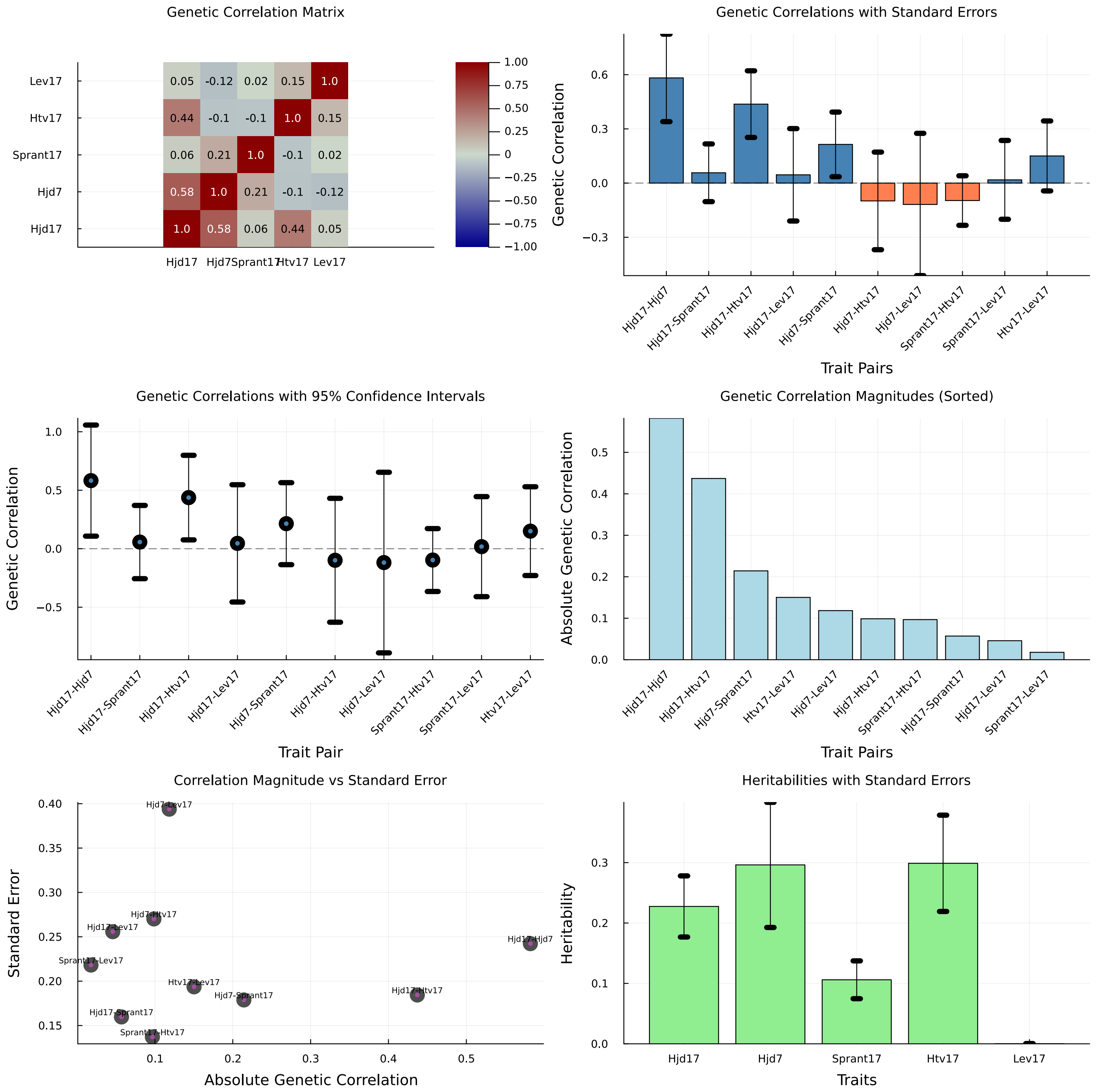


**Figure S5** Genetic correlations among growth and wood quality traits in the genotyped subset of Norway spruce. Results from multivariate mixed model analysis using genomic relationship matrix (G-matrix) for 1,218 genotyped individuals. (Top left) Genetic correlation matrix showing pairwise correlations with color intensity representing magnitude (blue = positive, red = negative). (Top right) Genetic correlations with standard errors, distinguishing positive (blue) from negative (red) correlations. (Middle left) Genetic correlations with 95% confidence intervals, revealing substantial uncertainty in most estimates. (Middle right) Absolute genetic correlation magnitudes sorted by strength. (Bottom left) Relationship between correlation magnitude and estimation precision (standard error). Most correlations cluster near zero with large standard errors, indicating limited power in the genotyped subset. (Bottom right) Narrow-sense heritabilities with standard errors for each trait. Traits: Hjd17 = dominant height at age 17, Hjd7 = dominant height at age 7, Sprant17 = number of ramicorns at age 17, Htv17 = total volume at age 17, Lev17 = survival at age 17. Compared to the full population (n=5,525), the genotyped subset shows substantially weaker correlations (e.g., Hjd17-Hjd7: 0.58 vs 0.90) and higher uncertainty, reflecting reduced statistical power from smaller sample size and potential sampling effects.

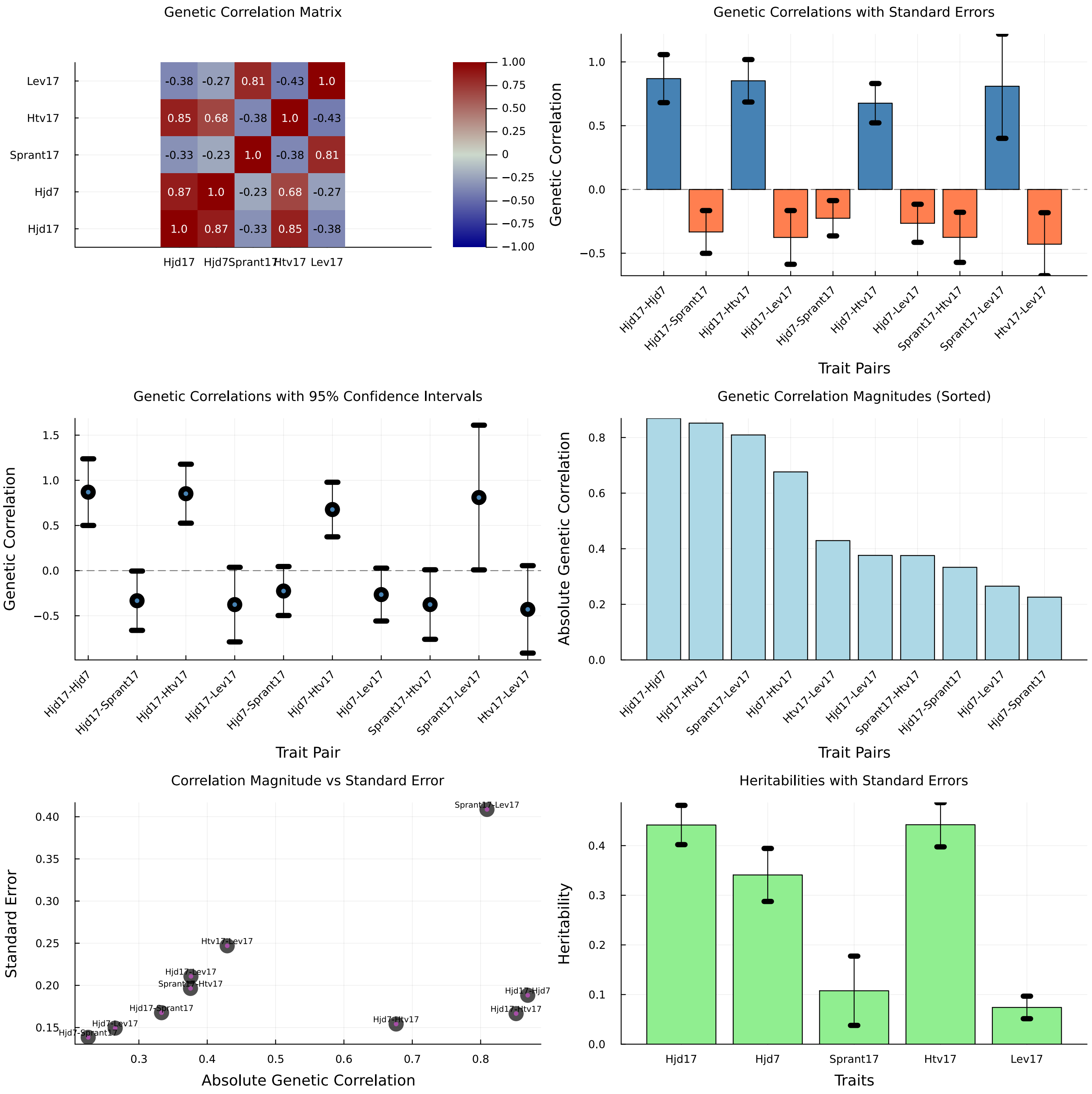


**Figure S6** Genetic correlations among growth and wood quality traits in Norway spruce. Results from multivariate mixed model analysis of 5,525 individuals from both field trials using blended A and G relationship matrices (e.g. H matrix). (Top left) Genetic correlation matrix showing pairwise correlations with color intensity representing magnitude (blue = positive, red = negative). (Top right) Genetic correlations with standard errors, distinguishing positive (blue) from negative (red) correlations. (Middle left) Genetic correlations with 95% confidence intervals, highlighting uncertainty in estimates. (Middle right) Absolute genetic correlation magnitudes sorted by strength, revealing trait pairs with strongest genetic associations. (Bottom left) Relationship between correlation magnitude and estimation precision (standard error), with trait pairs labeled. Strong correlations (e.g., Hjd17-Hjd7, Hjd17-Htv17) show lower uncertainty than moderate correlations (e.g., Sprant17-Lev17). (Bottom right) Narrow-sense heritabilities with standard errors for each trait. Traits: Hjd17 = dominant height at age 17, Hjd7 = dominant height at age 7, Sprant17 = ramicorn number at age 17, Htv17 = total volume at age 17, Lev17 = survival at age 17. Strong positive correlations exist among growth traits (Hjd17-Hjd7: 0.90, Hjd17-Htv17: 0.85), while ramicorn number shows negative genetic correlations with growth traits, indicating favorable breeding prospects for simultaneous improvement of growth and wood quality.

##### Benchmark OCS solver comparisons

As the JuMP suite of optimization tools contains a variety of optimizers, we were interested in comparing the performance on the Norway spruce breeding population to select the best performing method for in-depth uncertainty and robustness analyses. We compared four optimizers: Ipopt, ECOS, SCS and COSMO, all based on different techniques (Table S5). All methods successfully converged to optimal solutions, but differed substantially in computational efficiency.

The choice of relationship matrix significantly affected computational requirements. For the Norway spruce subset ($n = 1{,}218$), genomic relationship matrices (**G**) required 4.48–75.71 seconds, while the corresponding additive relationship matrices (**A**) were solved in 1.17–4.08 seconds. This 4–18 fold difference highlights the computational complexity introduced by non-sparse genomic relationships.

Performance differences became more pronounced with larger population sizes. For the **A** matrix with 5,525 individuals, COSMO (26.62 ± 0.45 s) and SCS (26.78 ± 2.76 s) maintained excellent performance, while Ipopt (18.11 ± 0.38 s) showed the fastest execution time. ECOS required substantially more time (601.00 ± 5.47 s), representing a 20–30 fold increase compared to the best-performing methods.

The **H** matrix ($n = 5{,}525$) presented the greatest computational challenge for all methods. COSMO achieved the fastest solution time (431.06 ± 9.63 s), followed by SCS (472.03 ± 12.48 s). Ipopt's performance degraded significantly (818.89 ± 15.93 s), while ECOS became impractical for routine use (5,086.92 ± 34.61 s).

These results demonstrate that algorithm selection should consider both population size and matrix structure. For routine applications involving large genomic datasets, COSMO and SCS provide the most reliable performance, while Ipopt remains competitive for smaller problems and pedigree-based relationship matrices. For the remainder of the paper, all analyses were performed using the COSMO optimizer.

#### Robustness score results

##### Norway spruce

For the genotyped Norway spruce subpopulation ($n = 1{,}218$), MAP-OCS selected 154 individuals at $\Theta = 0.02$ ($G_{\text{MAP}} = 1.46$). Applying the dynamic bottom-quartile threshold identified 25 high-risk selections (mean $S_i = 0.013 \pm 0.008$) whose robustness scores were 226% lower than the remaining MAP-selected individuals (mean $S_i = 0.044 \pm 0.027$; Fig. S7). Notably, three of the top-25 MAP contributors were classified as high-risk, including one in the top-10 — illustrating that high contribution under MAP EBVs does not guarantee stability under posterior uncertainty.

Re-running OCS after excluding these 25 individuals improved mean robustness by 16.5% at a genetic gain cost of only 3.0% ($G_{\text{constrained}} = 1.42$), with 128 individuals (81.1%) shared between the two solutions (Table S6). Excluding high-risk individuals increased contribution concentration among the retained selections: the Gini coefficient rose by +5.6% and top-10% concentration increased by +12.9%, reflecting the structural limitation of post-hoc exclusion discussed in the main manuscript.

**Table S5** Available reference methods used to compare the performance of the optimization implementations. Time in bold indicates the fastest method per comparison.

| Library | Method | Reference | Pedigree | NRM | Size | Time [s] |
|---|---|---|---|---|---|---|
| Ipopt | Interior point | Wächter and Biegler (2006) | Norway spruce | **G** | 1,218 | 6.40 (0.23) |
| | | | | **A** | 1,218 | **1.17 (0.03)** |
| | | | | **A** | 5,525 | **18.11 (0.38)** |
| | | | | **H** | 5,525 | 818.89 (15.93) |
| ECOS | Embedded conic solver | Domahidi *et al.* (2013) | Norway spruce | **G** | 1,218 | 75.71 (2.54) |
| | | | | **A** | 1,218 | 4.08 (0.09) |
| | | | | **A** | 5,525 | 601.00 (5.47) |
| | | | | **H** | 5,525 | 5086.92 (34.61) |
| COSMO | Conic operator splitting | Garstka *et al.* (2019) | Norway spruce | **G** | 1,218 | **4.48 (0.02)** |
| | | | | **A** | 1,218 | 1.19 (0.05) |
| | | | | **A** | 5,525 | 26.62 (0.45) |
| | | | | **H** | 5,525 | **431.06 (9.63)** |
| SCS | Splitting cone solver | JuMP Dev. (2024) | Norway spruce | **G** | 1,218 | 4.49 (0.07) |
| | | | | **A** | 1,218 | 1.81 (0.12) |
| | | | | **A** | 5,525 | 26.78 (2.76) |
| | | | | **H** | 5,525 | 472.03 (12.48) |

Values in parentheses are standard deviations. **G**: genomic relationship matrix; **A**: pedigree relationship matrix; **H**: blended genomic and pedigree relationship matrix. NRM: numerator relationship matrix type.

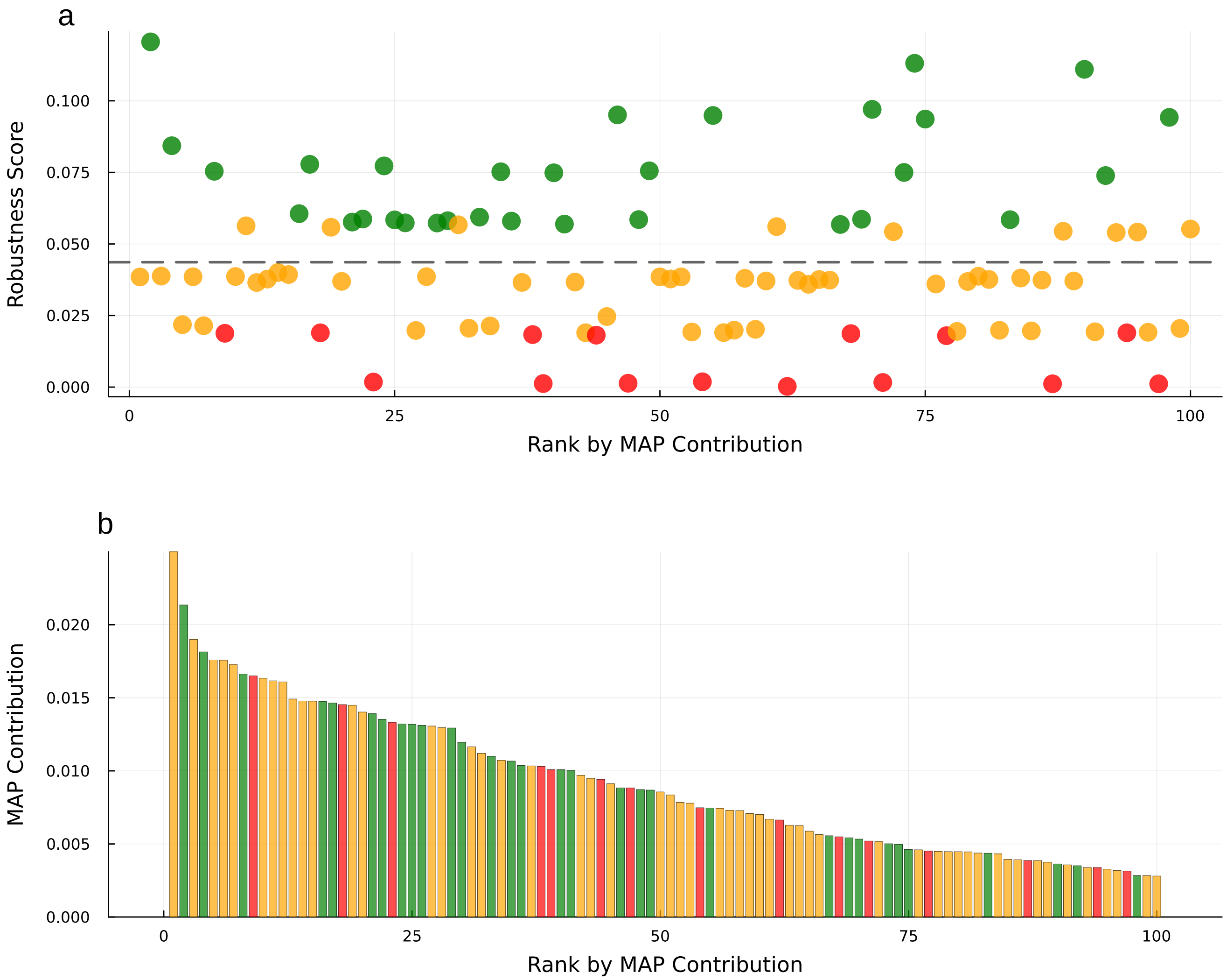


**Figure S7 Risk assessment of MAP-OCS selections based on MCMC robustness analysis in the Norway spruce genotyped subpopulation. a** Relationship between standardised breeding values and MCMC robustness scores for the top 100 MAP-OCS selections, coloured by risk category: green = Low Risk (score $\geq 1.5\times$ median), orange = Moderate Risk, red = High Risk (score $\leq 0.85\times$ median). Higher scores indicate individuals whose removal causes greater gain loss across MCMC iterations. **b** MAP contribution values for the same individuals ranked by contribution magnitude. Some high-contributing individuals carry low robustness scores, indicating suboptimal selection decisions when EBV uncertainty is considered.

**Table S6** Dual-metric summary for Norway spruce (*Picea abies*; $n$ = 1,218; $\Theta = 0.02$): MAP-OCS vs. constrained OCS after exclusion of bottom-quartile robustness selections.

| Method | $n_{\text{sel}}$ | $\bar{G}$ | Mean $S_i$ | Gini | $\text{Conc}_{10}$ (%) | Gain loss (%) |
|---|---|---|---|---|---|---|
| MAP-OCS | 154 | 1.46 | $0.036 \pm 0.027$ | — | — | — |
| Constrained OCS | 128 | 1.42 | $0.042 \pm 0.024$ | — | — | −3.0 |

Gini and $\text{Conc}_{10}$ absolute values not available in current output; percentage changes reported in main text (+5.6% and +12.9% respectively).

#### CVaR-OCS results across coancestry constraints (QTL-MAS 2010)

Full CVaR-OCS results across all three coancestry constraints ($\Theta \in \{0.02, 0.03, 0.05\}$) for the QTL-MAS 2010 simulation dataset are presented in Table S7. The recommended operating point for each $\Theta$ is identified based on the trade-off between expected gain and tail-risk protection.

#### MCMC-OCS overlap metrics by relationship matrix (Norway spruce)

Table S8 summarises the overlap between MAP-OCS and MCMC-OCS solutions across all relationship matrices for Norway spruce. Key results for the primary genomic analysis (**G**-matrix, $n = 1{,}218$) are discussed in the main text.

#### Computational cost of CVaR-OCS and robustness score calculations

To address potential concerns about computational scalability, we benchmarked the wall-clock time of MAP-OCS and CVaR-OCS on all datasets used in this study (Table S9). All timings were obtained on a standard desktop workstation running Windows 11 with an Intel Core i7 processor. Analyses were implemented in Julia using the COSMO solver (Garstka *et al.* 2019) via JuMP. To avoid inflation from Julia's just-in-time (JIT) compilation, one untimed warm-up solve was performed for each method before recording the $N = 10$ replicate runs reported here.

A single CVaR-OCS solve required approximately 22× longer than MAP-OCS for the Norway spruce G-matrix subset ($n = 1{,}218$; $84.0 \pm 4.4$ s vs. $3.9 \pm 0.9$ s), and approximately 11× longer for the full QTLMAS dataset ($n = 3{,}226$; $1{,}718 \pm 52$ s vs. $155 \pm 20$ s). The overhead reflects the $l = 1{,}000$ auxiliary variables and $l$ shortfall constraints added to the quadratic programme by the CVaR formulation, and scales with both $n$ and $l$. Absolute solve times for the smaller datasets remained well within practical limits; the larger datasets ($n > 3{,}000$) require more careful scheduling but remain feasible given that CVaR-OCS is a one-time-per-generation operation. Note that the QTLMAS benchmark used the full genomic relationship matrix across all generations ($n = 3{,}226$), whereas operational use would typically be restricted to the current selection candidates.

The parameter frontier sweep across all $|\mathcal{A}| \times |\mathcal{M}| = 3 \times 9 = 27$ $(\alpha, \mu)$ combinations is a one-time analysis cost; once a preferred operating point is identified it need not be repeated each generation. Its cost is fully explained by $27 \times t_{\text{CVaR}}$, where $t_{\text{CVaR}}$ is the mean single-solve time (Table S9).

Robustness score computation requires one MAP-OCS solve per candidate per MCMC posterior draw used in the leave-one-out loop. For the Norway spruce G-matrix subset ($n = 1{,}218$) this amounted to $n_{\text{sel}} \times n_{\text{MCMC}} = 88 \times 200 = 17{,}600$ MAP-OCS calls (approximately 1,133 min sequentially, or ∼57 min on 20 cores); for the full Norway spruce population ($n = 5{,}525$) the corresponding figure was $41 \times 200 = 8{,}200$ calls (∼22,079 min sequentially, ∼1,104 min on 20 cores). Because each candidate's score is computed independently, the loop is embarrassingly parallel; reported estimates assume sequential execution, with a 20-core parallel estimate included for reference.

**Table S7 Full CVaR-OCS results for the QTL-MAS 2010 simulation study** ($n = 900$ candidates; $l = 5{,}000$ MCMC scenarios) across all three coancestry constraints ($\Theta \in \{0.02, 0.03, 0.05\}$). CVaR-OCS is reported at the recommended operating point for each $\Theta$: $(\alpha^*, \mu^*) = (0.95, 1.5)$ at $\Theta = 0.02$; $(0.90, 1.8)$ at $\Theta = 0.03$; $(0.90, 1.5)$ at $\Theta = 0.05$. $E[\text{gain}]$: in-sample expected genetic gain[a]; Realised gain: $\mathbf{c}^\top\mathbf{t}$ evaluated against true breeding values (TBVs); $\text{CVaR}_{95}$: expected gain in the worst 5% of MCMC scenarios; $n_{\text{sel}}$: number of selected parents (sires S, dams D); Jaccard: set overlap with oracle. $\Delta(\%)$ is relative to MAP-OCS(EBV) within each $\Theta$ column. Results at $\Theta = 0.03$ are also presented in the main text (Table 2 therein).

| | | $\Theta = 0.02$ | | $\Theta = 0.03$ | | $\Theta = 0.05$ | |
|---|---|---|---|---|---|---|---|
| Metric | Solution | Value | $\Delta(\%)$ | Value | $\Delta(\%)$ | Value | $\Delta(\%)$ |
| | Oracle | 1.599 | — | 1.725 | — | 1.861 | — |
| $E[\text{gain}]$ | MAP-OCS(EBV) | 1.440 | — | 1.595 | — | 1.726 | — |
| | CVaR-OCS | 1.440 | +0.0 | 1.596 | +0.1 | 1.730 | +0.2 |
| Realised gain | MAP-OCS(EBV) | 1.195 | — | 1.342 | — | 1.527 | — |
| | CVaR-OCS | 1.187 | −0.7 | 1.325 | −1.2 | 1.506 | −1.4 |
| $\text{CVaR}_{95}$ | MAP-OCS(EBV) | 1.278 | — | 1.400 | — | 1.474 | — |
| | CVaR-OCS | 1.286 | +0.6 | 1.437 | +**2.7** | 1.523 | +**3.4** |
| $\text{VaR}_{95}$ | MAP-OCS(EBV) | 1.311 | — | 1.440 | — | 1.524 | — |
| | CVaR-OCS | 1.317 | +0.4 | 1.469 | +2.1 | 1.537 | +0.8 |
| | Oracle | 44 (19, 25) | — | 31 (15, 16) | — | 19 (9, 10) | — |
| $n_{\text{sel}}$ (S, D) | MAP-OCS(EBV) | 53 (28, 25) | — | 29 (17, 12) | — | 20 (12, 8) | — |
| | CVaR-OCS | 59 (33, 26) | +11.3 | 35 (20, 15) | +20.7 | 24 (13, 11) | +20.0 |
| Gini | MAP-OCS(EBV) | 0.467 | — | 0.347 | — | 0.407 | — |
| | CVaR-OCS | 0.475 | +1.7 | 0.407 | +17.3 | 0.460 | +13.0 |
| Jaccard vs oracle | MAP-OCS(EBV) | 0.228 | — | 0.176 | — | 0.258 | — |
| | CVaR-OCS | 0.226 | −0.9 | 0.196 | +11.4 | 0.229 | −11.2 |
| Shared with oracle | MAP-OCS(EBV) | 18 | — | 9 | — | 8 | — |
| | CVaR-OCS | 19 | +5.6 | 11 | +22.2 | 8 | +0.0 |

[a] MAP-OCS(EBV) $E[\text{gain}]$ is optimised against the posterior-mean EBV vector and overestimates realised gain by 20.5%, 18.8%, and 13.1% at $\Theta = 0.02$, 0.03, and 0.05 respectively, due to EBV uncertainty. CVaR-OCS $E[\text{gain}]$ is jointly optimised over all $l$ scenarios and carries no such bias. Bold $\Delta(\%)$ values indicate the recommended operating point per $\Theta$.

**Table S8** MCMC-OCS uncertainty analysis: summary statistics of primary overlap metrics for Norway spruce by relationship matrix. Key numbers for the primary genomic analysis (**G**-matrix, $n = 1{,}218$) are reported in the main text.

| Species | Matrix | Pop Size | $\Theta$ | Overlap Count | Jaccard Sim |
|---|---|---|---|---|---|
| Norway Spruce | A | 1,218 | 0.02 | 24.9 (4.3) | 0.143 (0.028) |
| Norway Spruce | G | 1,218 | 0.02 | 26.6 (4.8) | 0.154 (0.032) |
| Norway Spruce | A | 5,525 | 0.02 | 10.4 (4.0) | 0.056 (0.023) |
| Norway Spruce | H | 5,525 | 0.02 | 14.1 (3.6) | 0.076 (0.021) |

Values in parentheses are standard deviations across MCMC iterations. **G**: genomic relationships; **A**: pedigree relationships; **H**: blended genomic and pedigree relationships; $\Theta$: group coancestry constraint. Overlap Count is the mean number of individuals selected in both the MAP-OCS and MCMC-OCS solutions out of 100 total selections per iteration.

**Table S9** Wall-clock benchmarks for MAP-OCS and CVaR-OCS across datasets. Timed methods report mean ± SD over $N = 10$ replicate solves after one untimed warm-up call. Sweep and robustness score times are extrapolated (footnotes *b–c*; see Section ). $n$: genotyped candidates; $l$: MCMC posterior samples used as CVaR scenarios; $\Theta$: group coancestry constraint; $n_{\text{sel}}$: individuals selected by MAP-OCS; $n_{\text{MCMC}}$: posterior draws used per candidate in robustness score calculation.

| Dataset | Method | $n$ | Mean (s) ± SD | ×MAP | Sequential (min) | 20-core (min)[a] |
|---|---|---|---|---|---|---|
| *Norway spruce G-matrix* ($\Theta = 0.02$, $l = 1{,}000$, $n_{MCMC} = 200$) | | | | | | |
| Spruce ($n = 1{,}218$) | MAP-OCS | 1,218 | $3.9 \pm 0.9$ | ref | $< 0.1$ | — |
| | CVaR-OCS (single) | 1,218 | $84.0 \pm 4.4$ | ×21.8 | 1.4 | — |
| | CVaR-OCS (sweep)[b] | 1,218 | $27 \times 84.0 = 2{,}269$ s | | 37.8 | — |
| | Robustness scores[c] | 1,218 | $88 \times 200 \times 3.9 = 67{,}989$ s | | 1,133 | 57 |
| Spruce ($n = 5{,}525$) | MAP-OCS | 5,525 | $161.6 \pm 23.0$ | ref | 2.7 | — |
| | CVaR-OCS (single) | 5,525 | $2{,}724 \pm 41$ | ×16.9 | 45.4 | — |
| | CVaR-OCS (sweep)[b] | 5,525 | $27 \times 2{,}724 = 73{,}553$ s | | 1,226 | — |
| | Robustness scores[c] | 5,525 | $41 \times 200 \times 161.6 = 1{,}325{,}000$ s | | 22,079 | 1,104 |
| *QTLMAS 2010, all generations* ($\Theta = 0.05$, $l = 1{,}000$, $n = 3{,}226$, $n_{MCMC} = 200$) | | | | | | |
| QTLMAS ($n = 3{,}226$) | MAP-OCS | 3,226 | $154.7 \pm 19.7$ | ref | 2.6 | — |
| | CVaR-OCS (single) | 3,226 | $1{,}718 \pm 52$ | ×11.1 | 28.6 | — |
| | CVaR-OCS (sweep)[b] | 3,226 | $27 \times 1{,}718 = 46{,}376$ s | | 773 | — |
| | Robustness scores[c] | 3,226 | $13 \times 200 \times 154.7 = 402{,}174$ s | | 6,703 | 335 |

[a] Indicative parallel estimate assuming embarrassingly parallel execution over candidates (20 cores); actual speedup depends on hardware and memory bandwidth.

[b] Extrapolated as $27 \times \bar{t}_{\text{CVaR,single}}$; the sweep is a one-time parameter selection cost, not repeated each generation.

[c] Extrapolated as $n_{\text{sel}} \times n_{\text{MCMC}} \times \bar{t}_{\text{MAP}}$; independent over candidates and therefore embarrassingly parallel.
